# Winding-Up of Fibrin Fibers as a Novel Mechanism of Platelet-Mediated Fiber Compaction

**DOI:** 10.64898/2026.02.15.705975

**Authors:** Alexei Grichine, Tatiana Kovalenko, Florence Appaix, Anne-Sophie Ribba, Anita Eckly, Jean-Yves Rinckel, Mikhail Panteleev, Laurence Lafanechère, Karin Sadoul

## Abstract

This study reveals a previously unrecognized mechanism by which platelets retract and compact fibrin fibers. Using a newly developed 2D fiber-retraction assay, we observed an initial “gearwheel” pattern of actin–myosin organization in spread platelets with associated, extracellular fibrin patches that appear to form an initiation complex for fibrin fiber attachment and rearrangement. The final outcome of this process results in two morphologically different platelet populations. Spread platelets with coiled fibers around and above their pseudo-nucleus. Other platelets are only partially spread on the glass surface and are surrounded by tightly packed fibrin fibers around bulbous protrusions (“bulbs”), mirroring the architecture of platelets and adjacent fibers within a retracted clot. Thus, the observed compaction process might also take place during clot retraction in order to reduce clot volume, stiffen the clot and enhance wound repair. Apart from pulling on fibers like on a rope, platelets actively wrap fibrin fibers into compact structures, similar to balls of wool. Besides DNA packaging, this represents a new example of a natural fiber compaction mechanism. Using a combination of 3D clot-retraction and 2D fiber-retraction assays, expansion and electron microscopy, live imaging and mathematical modeling, we show that platelets use an actomyosin-driven motion to gather and loop fibrin fibers around the base of bulbous protrusions. These bulbs form when a platelet becomes trapped between fibrin fibers during 3D clot retraction or 2D fiber-retraction assays. These findings complement and extend earlier models of platelet-mediated fibrin fiber retractions, offering new insight into how platelets mechanically organize fibrin fibers.

## Introduction

Platelets are absolutely essential in case of vessel injury. They are not only necessary to initiate hemostasis but also to retract the forming clot thereby pulling together the edges of the ruptured vessel and flattening the clot for improved blood flow. Different approaches to visualize fibrin fibers in whole blood or plasma clots have shown that fibrin fibers represent the main mechanical and structural framework of clots,^1,2^ occupying a substantial portion of the clot volume. Previous studies have shown that platelets are able to retract an unconstrained plasma clot to a tiny volume.^3-6^ The extent and rate of clot retraction are influenced by several factors, including platelet and red blood cell counts, fibrin and thrombin concentrations, and the geometric context of clot formation.^7^ However, platelets are the primary cellular agents driving such extensive clot compaction, although erythrocytes also contribute.^8^ How do platelets accomplish this tremendous task? Two key studies offer insight: While Fletcher’s lab demonstrated that platelets generate contractile forces comparable to muscle cells,^9^ Weisel’s group provided mechanistic insight, showing that platelets extend filopodia to bind and pull on fibrin fibers, much like tugging on ropes^10^ causing localized fiber accumulation.

However, pulling alone may not fully account for the dramatic reduction in clot volume. We hypothesized that efficient clot retraction requires substantial densification of the fibrin fiber network. Using expansion microscopy, we observed that fibrin fibers encircle the central region of platelets within a constrained clot, forming a cage-like structure. Using a “2D fiber-retraction assay” we found that fibrin patches are initially arranged in a ring-like pattern in close proximity to the circular organization of the actin cytoskeleton in spread platelets. Over time, a myosin-dependent mechanism drives the accumulation of coiled fibrin fibers above the plasma membrane of fully spread platelets and their compaction around platelet bulbs in partially adherent platelets.

To explore the mechanistic aspects of the platelet mediated fiber compaction we used computational modeling and live-cell video microscopy. Our results suggest that actomyosin-driven swirling within spread platelets on a 2D surface or within individual bulbs of platelets in the 3D clot space may underlie the observed fibrin coiling and compaction.

## Results

### Platelets become labeled by fluorescent fibrin fibers during clot retraction

We first estimated the capacity of platelets to retract diluted plasma clots (12% plasma) under unconstrained conditions (Fig. 1A). Under these conditions, 4×10^7^ platelets are able to retract the clot volume to approximately 1.5% of its initial volume (400 μl) within 60 min. Even platelets kept overnight at room temperature before the retraction assay were still retracting the clot to the same extent, however after a longer lag phase. We then used live imaging to follow the reorganization of fibrin fibers during unconstrained clot retraction. Platelet rich plasma (PRP) was spiked with fibrinogen-Alexa 488 before thrombin induced clot formation and image acquisition. Very rapidly one can observe the formation of fluorescent fibrin nodes during the retraction process. Since under the assay conditions the clot is composed only of platelets and fibrin fibers, we hypothesized that the nodes may correspond to platelets surrounded by fibrin fibers (Fig. 1B and video 1). To test this hypothesis, we performed a time course analysis of clot retraction in presence of fluorescent fibrinogen under constrained conditions preventing complete retraction. This approach allowed us to stain and visualize individual platelets in a clot even after prolonged retraction times. Hemostatic clots that form in vivo following vessel injury are also likely to experience mechanical constraints, as they remain anchored to the damaged vessel walls while spanning the breach. Retracted clots were fixed and stained for the αIIb integrin subunit to detect the localization of platelets within the clot. At the position of each platelet, we consistently see colocalization of a fibrin node at all time points examined (Fig. 1C), confirming our hypothesis that the fibrin nodes observed in figure 1B form at platelet sites. Furthermore, the results show that the build-up of fibrin fibers around individual platelets occurs very rapidly, within 10 min. The fibrin fibers forming the nodes did not decorate the entire platelet surface but surrounded the center of platelets embedded in the clot. Please note, that platelets occupy a larger space after longer retraction times, but the fiber nodes persist even after 80 min of retraction.

**Figure 1:**
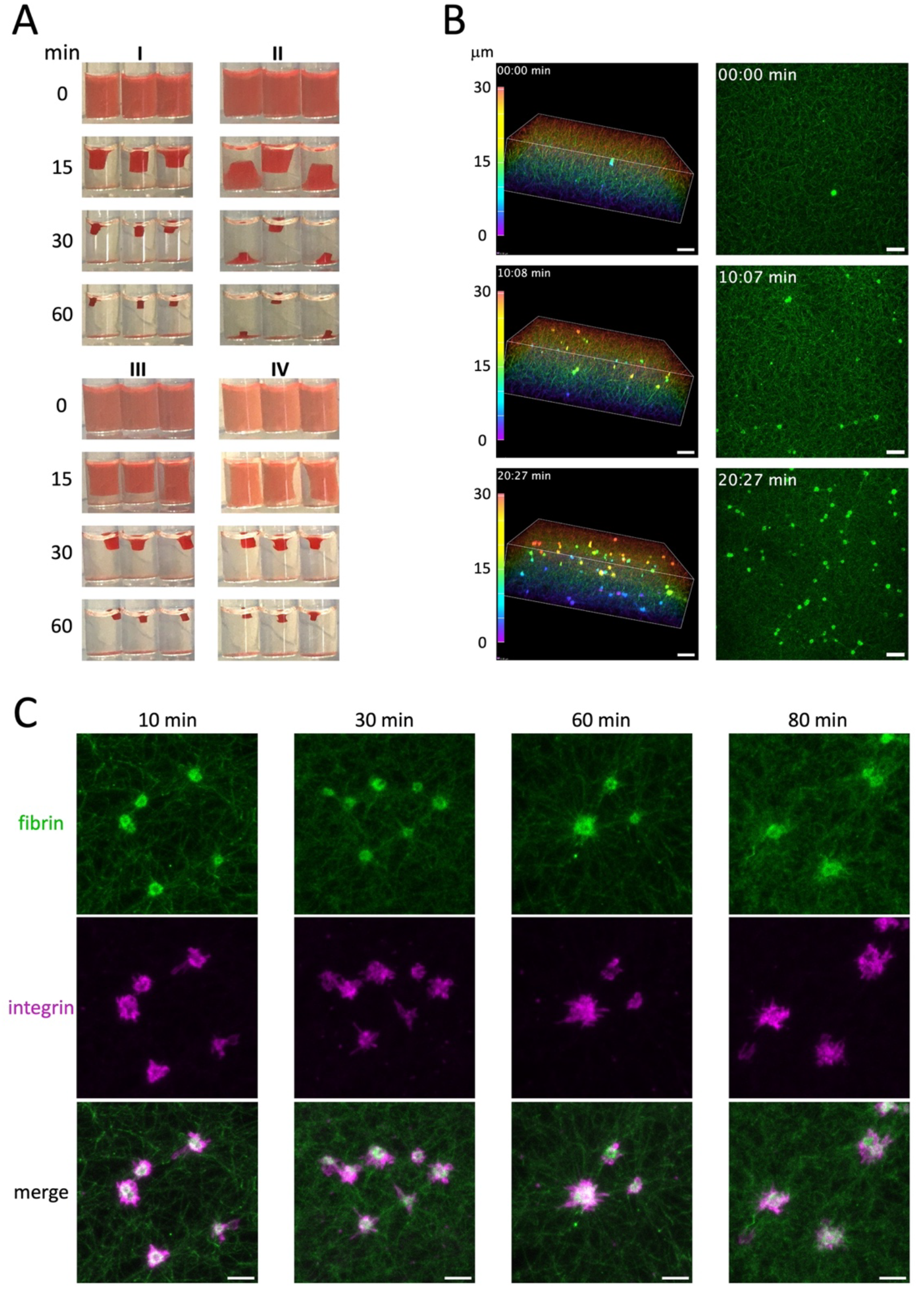
Platelets become labelled by fluorescent fibrin fibers during clot retraction. A)Time course of unconstrained clot retraction by platelets (4×10^7^ in 400 μl of 12% plasma in PBS) for a total of 60 minutes at room temperature. Experiments have been performed 6x using blood from different donors and typical examples of blood from two different donors are shown (experiments III and IV are repetitions of I and II the day after; although a longer lag-phase is observed, the final retraction volume is similar for the four experiments). B)Unconstrained clot retraction (1×10^8^ platelets per ml in 50% plasma/50% PBS) in presence of fibrinogen-Alexa 488. Image acquisition was started immediately after thrombin addition at a focal plane 100 μm above the bottom of the well and image stacks were collected (61 focal planes, step size 0.5 μm) for a time period of 20:27 min (100 frames with a time interval of 12.5 sec). See also associated video 1. The experiment has been performed twice using platelets of the same donor. Shown are the first, intermediate and last time points of a depth color-coded time-lapse video (left panel; scale bar 10 μm) and the maximum intensity projections (MIPs) of the same time points (right panel; scale bar 10 μm). C)Constrained clot retraction (1×10^8^ platelets per ml in 50% plasma/50% PBS, fibrinogen-Alexa 488 final concentration 12.5 μg/ml) between two holders. Clots were induced by addition of thrombin (2.5U/ml final), fixed at the indicated retraction times, embedded in gelatin, flash frozen and cryosections (14 μm) were stained for the σIIb integrin subunit (magenta; scale bar 5 μm). The time course was performed twice using PRP from two different donors (retraction assays were repeated, although not for all time points, more than eight times using blood from different donors, with consistent results). Image acquisition was performed using a wide-field epi fluorescence microscope (BX41; Olympus) equipped with a Plan 100x/1.25 NA oil objective, a camera (DP70; Olympus), and the acquisition software analySIS (Olympus).

To gain a better insight about the organization of fibrin fibers around platelets within a clot, we used 4x isotropic expansion microscopy. We found that fibrin fibers form a cage-like structure around the platelet center, clearly visible when scrolling through the image stacks (Fig. 2 and video 2 with compiled animations of image parts 2A-E). This pattern was consistently observed across all platelets in the clot and after different clot retraction times (15 min Fig. 2A-D and 4h Fig. 2E). Most platelets had formed large bulbs, thin filopodia, or both with fibrin fibers typically localized around the base of the bulbs or along filopodia. It remains unclear whether all fibrin segments reside on the external platelet membrane or whether some of them extend into channels of the open canalicular system (OCS). Some of these channels persist in activated platelets.^11^ Please note that straight fibers radiate from the platelet, indicating a high tension in the platelet clot environment. We want to emphasize that changes in temperature, or plasma/platelet concentrations change the kinetics of retraction but we always observe the cage-like fibrin fiber organization around individual platelets (not shown). We also investigated the organization of the actin cytoskeleton of platelets in a constrained clot and demonstrated the presence of radial actin filaments in each bulb, extending to the center of the platelet, where the cage like fibrin fiber organization is observed (Fig. 2E and video 2).

**Figure 2:**
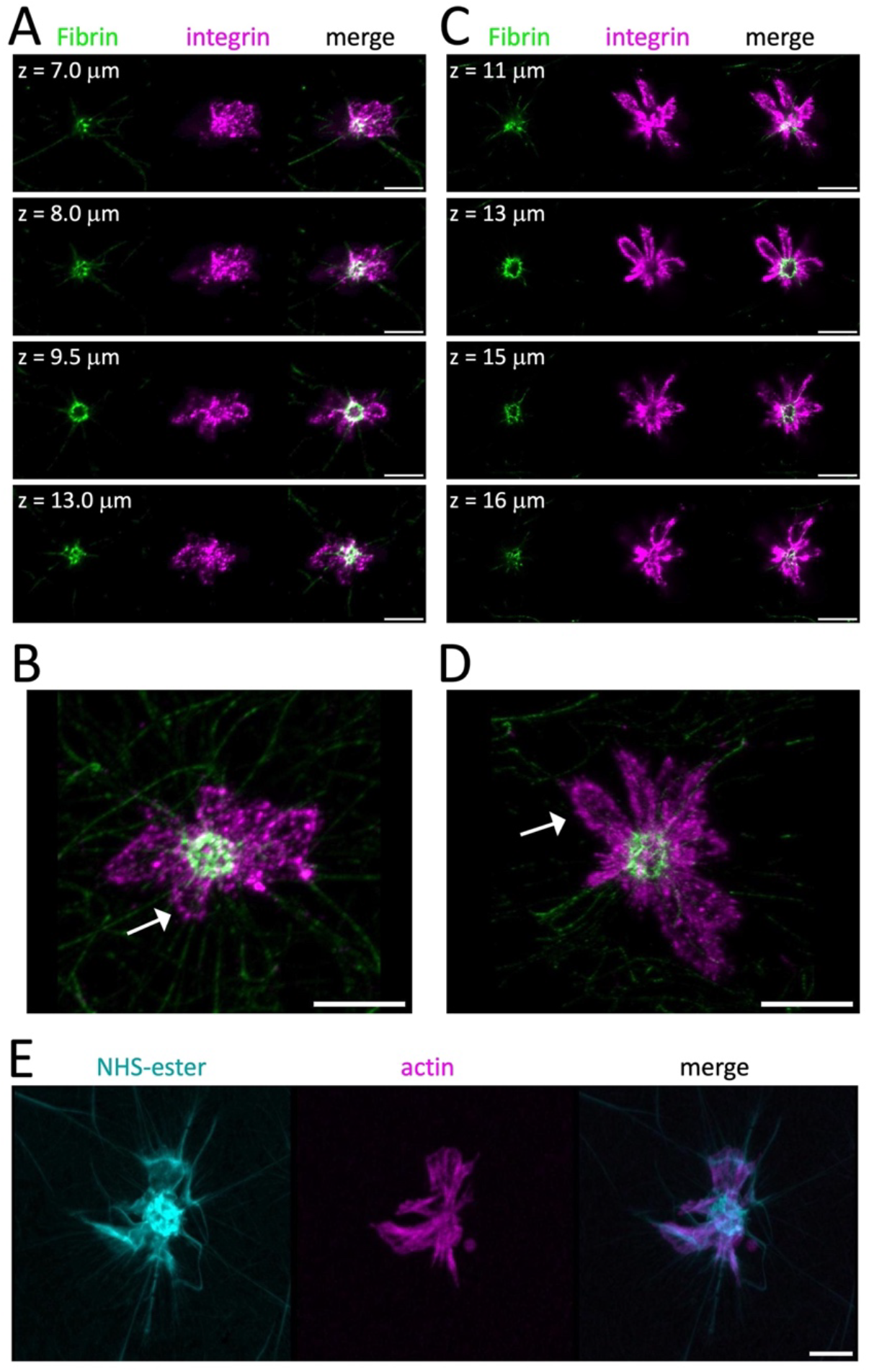
Fibrin fibers are organized in a “cage-like” fashion around platelets within a constrained clot. A-D: Two typical examples (out of n=39 acquisitions of four experiments using blood from different donors) of platelets in a constrained clot with attached fibrin fibers are shown (1×10^7^ platelets per ml PBS/50% plasma and fibrinogen-Alexa 488). Clot retraction was allowed to take place in an inoculation loop (see methods section) for 15 min before fixation and immunofluorescence staining of the σIIb integrin subunit (magenta). Samples were then processed for expansion (scale bars 10 μm = 2.5 μm after correction for expansion; indicated z-levels are not corrected for expansion; see also associated video 2 with compiled animations of image parts A-E). A) Four focal planes from a stack of images (z-level as indicated) showing a platelet in a clot (fibrin fibers in green, plasma membrane staining using an antibody against the σIIb integrin subunit in magenta and the merge). B) D reconstruction of the image stack shown in A (46 planes, step size 0.5 μm), a bulb is indicated by an arrow. C)Four focal planes show another typical platelet in a clot (z-level as indicated). D) 3D image reconstruction of the image stack shown in C (28 planes, step size 1 μm), a bulb is indicated by an arrow. E)3D image reconstruction of a platelet in a constrained clot (retraction in an inoculation loop for 4h) stained after expansion using the NHS-ester (cyan, for total protein stain) and an actin probe (magenta), demonstrating that radial actin fibers extending to the platelet center are present in each bulb (scale bar 10 μm = 2.5 μm after correction for expansion, see also video 2).

### Platelets, spread on a 2D surface, organize fibrin fibers above them

So far, our results demonstrate that platelets possess a remarkable capacity to precisely organize fibrin fibers around themselves when embedded in a clot. It is, however, difficult to find out how platelets integrated in the fiber network manage to do that. To investigate the mechanism by which platelets arrange the fibers around their center we sought to use a model system with reduced complexity. We, therefore, developed a “2D fiber-retraction assay”, which is a reductionist approach that does not replicate full clot retraction, but can provide insight into how platelets interact with and organize fibrin fibers. To this end, a diluted suspension of PRP (2.5×10^6^ platelets/ml, 4μl plasma/ml PBS) was incubated with fluorescent fibrinogen and thrombin for 15 min to induce platelet activation and fibrin polymerization. Platelets were then allowed to spread on a 2D surface for 30 min before fixation. Under these conditions, we observe four categories of spread platelets and associated fibrin fiber organizations (Fig. 3; for the criteria used to define the categories see Fig. 3C). The first category comprises platelets without fibrin fiber contact (28.3+2%). The second includes platelets with either a small fibrin dot at their center, which may represent a fibrin initiation complex, or a few fiber contacts above them (40+3%). A third category consists of platelets displaying an initiation complex with attached fibrin fibers and evidence of fiber buildup above the platelets (fiber accumulation 18.3+0.5%). A fourth category exhibits a high fiber densification above platelets (fiber compaction 13.6+0.9%) (Fig. 3A, left image; quantified in Fig. 3B). We then investigated, whether this fiber accumulation/compaction depends on myosin actions as it is the case for clot retraction. Myosin inhibition using blebbistatin strongly diminished the fiber buildup and densification above platelets (fiber accumulation 6.6+0.9% and fiber compaction 0.2+0.2%), while a higher percentage of platelets with fiber initiations or fiber contacts was observed (63.4+2%) (Fig. 3A, right image; quantified in Fig. 3B). For fibrin fiber bundling, staggered or crosslinked fibrin protofilaments no platelet actions are necessary as described previously.^12,13^ Since we see a clear difference between +/− blebbistatin conditions, demonstrating that platelet actions are required, we consider these alternative mechanisms to be unlikely.

**Figure 3:**
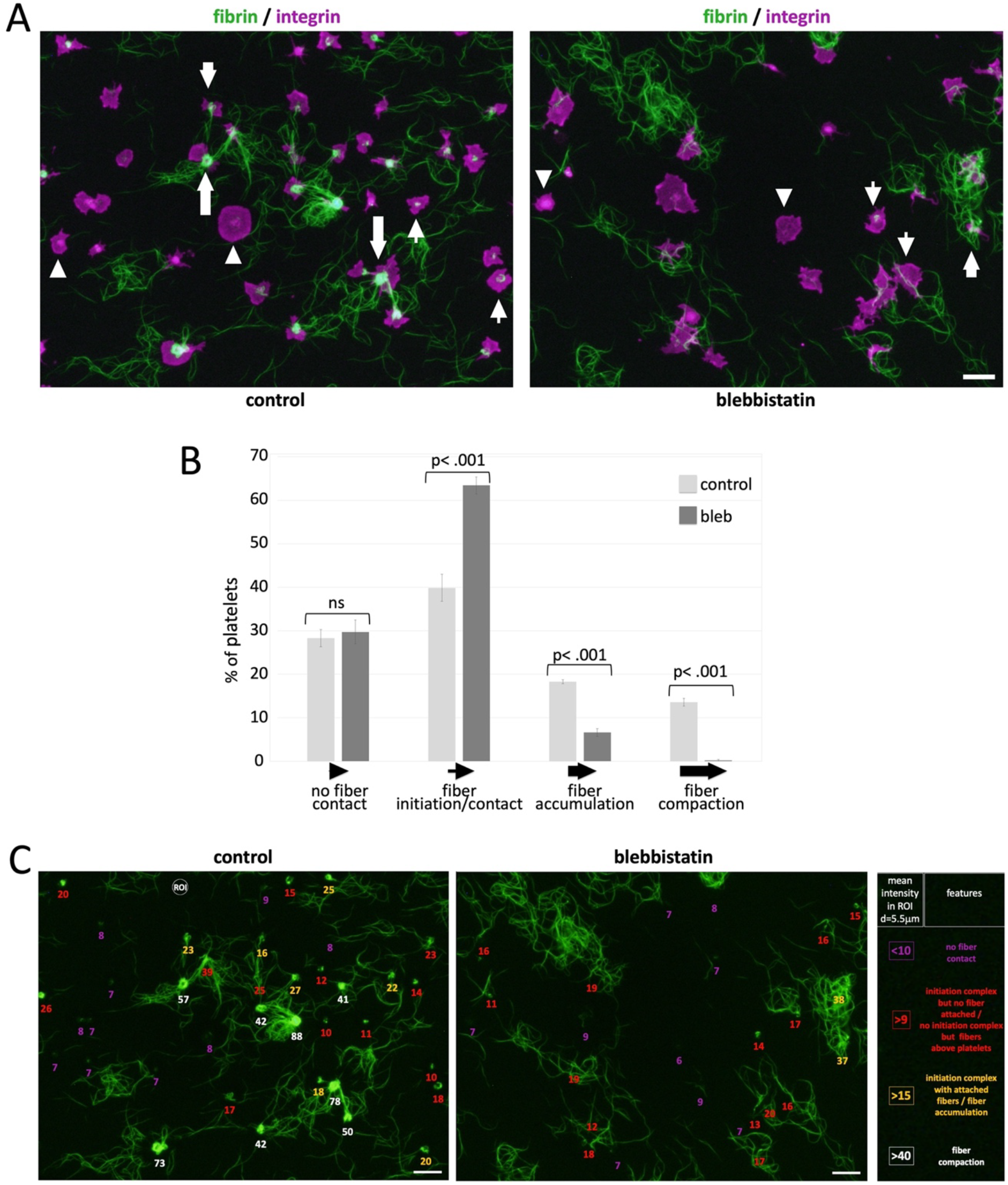
Spread platelets accumulate fibrin fibers above them in a myosin dependent way. A)Shown is a 2D fiber-retraction assay in the presence of DMSO (control) or 50 μM blebbistatin for myosin inhibition. After a 30 min preincubation of platelets (4 μl plasma per ml PBS, see methods section) with or without blebbistatin, thrombin is added and the assay is continued as described in the methods section. Scale bar 10 μm, arrowheads indicate platelets without fiber contact, thin arrows show platelets with a fiber initiation complex or with some fiber contacts. Larger arrows point to platelets with fiber accumulations and even larger arrows to platelets with compacted fibers (see legend below bar graph in B and the detailed features used to classify the different categories described in part C). B)Quantification of the percentage of platelets in the different categories using image acquisitions of four experiments using blood from different donors (for control conditions a total of 636 platelets were counted and for blebbistatin conditions a total of 524 platelets). Shown are means + sem, evaluation of significance using the Student’s t-test. C)Criteria used to classify the platelets and associated fibers shown in A. The green fluorescent fiber images shown in A were used to measure the mean fluorescence intensity of fibers present within a region of interest (ROI; diameter = 5.5 µm) at platelet sites. The measured values are shown below each platelet. The intensity values, together with the associated features (listed on the right), were used to define the four observed platelet categories.

Then, we used the 2D fiber-retraction assay followed by expansion microscopy to investigate in more detail how the fibers around platelets are organized. Platelets were stained either for the αIIb integrin subunit or for myosin. We observed platelets fully spread on the glass surface with coiled fibrin fibers above them (category “fiber accumulation” in fig. 3A). These fibers are frequently wrapped around a bulbous protrusion in the middle of the spread platelet, often referred to as pseudo-nucleus in the literature (Fig. 4A-E and video 3 with the compiled animations of image parts 4A-E). In this situation the cage-like fiber organization is disrupted.

**Figure 4:**
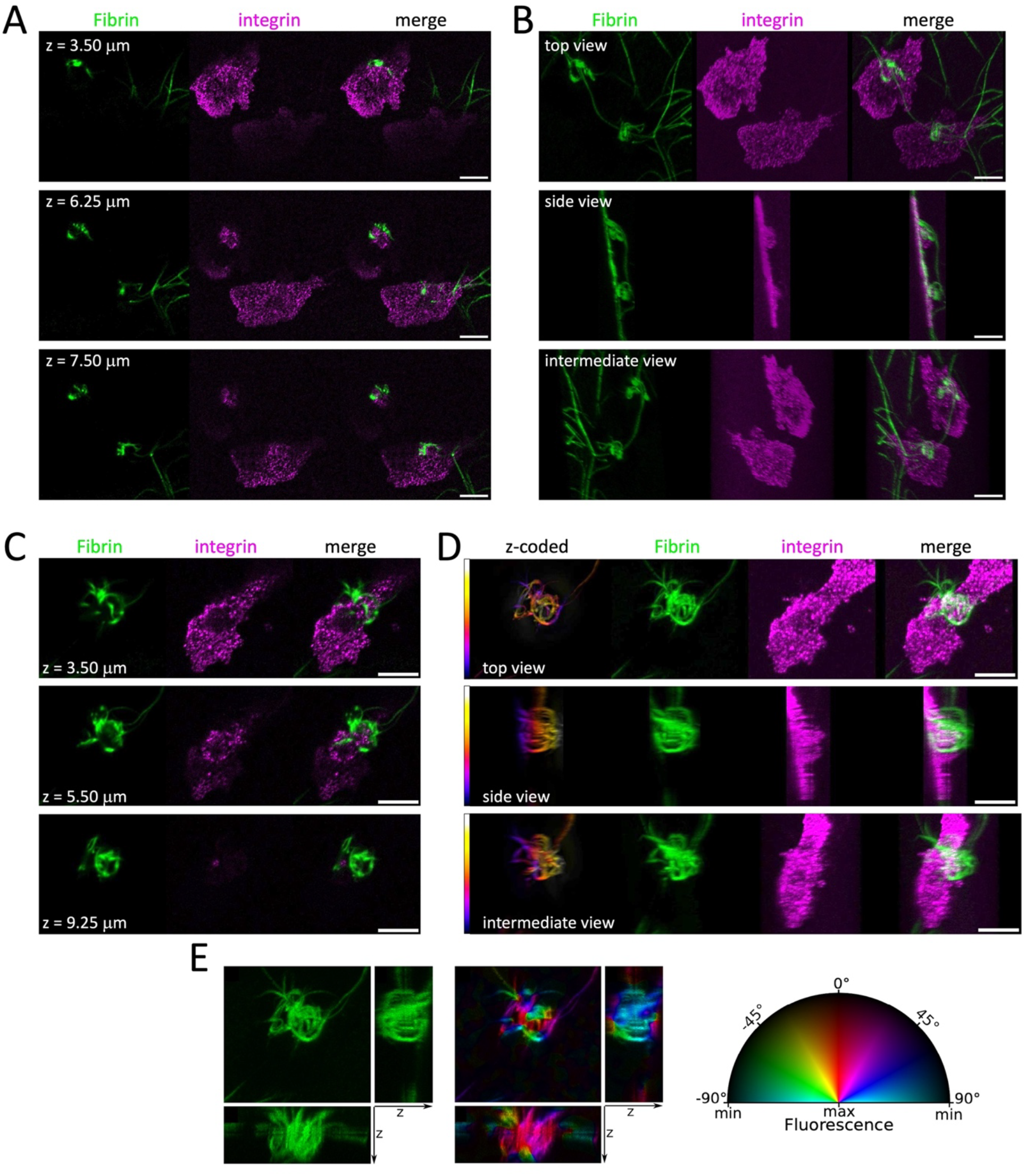
Coiled-up fibrin fibers above spread platelets. Two representative examples (out of n=18 acquisitions from three independent experiments using blood from different donors) of fibrin fiber accumulations above spread platelets in the 2D fiber-retraction assay (4 μl plasma per ml PBS, see methods section) are shown (see also video 3 with compiled animations of image parts A-D). Samples were stained for the αIIb integrin subunit and expanded (scale bars 10 μm = 2.5 μm after correction for expansion, indicated z-levels are not corrected for expansion). A) Three different focal planes (z-levels as indicated) from an image stack showing two spread platelets and attached fibrin fibers. B)3D reconstruction of the image stack shown in A (51 planes, step size 0.25 μm), represented are three different view angles. C)Three different focal planes from an image stack illustrating a spread platelet and attached, coiled-up fibrin fibers. D)3D reconstruction of the image stack shown in C (50 planes, step size 0.25 μm), represented are three different view angles. The fibrin fibers are additionally displayed as depth-color coded (Fire LUT) reconstructions to highlight the straight fiber parts along the z-axis. E)Same fiber image stack as in C and D. Left panels are maximum intensity projections of the green fibrin fibers along the z, x and y axes. Right panels are the same projections color coded in the HSB space (Hue, Saturation, Brightness) according to the orientation of individual fiber segments. The fibers are well aligned along the z-axis, while they crisscross horizontally.

However, in addition to these spread platelets with coiled fibrin fibers above them, we also observed platelets which form bulbs and filopodia similar to platelets within a clot. These platelets were only partially attached to the glass surface and a substantial fiber accumulation and densification was observed around the platelet center at the base of the bulbs (category “fiber compaction” in fig. 3). This fiber organization appeared similar to the cage-like structure formed by fibers around the center of platelets embedded in a clot. Nevertheless, under these 2D conditions, more densely compacted masses of fibers are observed around each bulb than under the 3D conditions of a clot. It is possible that several layers of fibers are tightly compacted around the base of the platelet bulbs, much like individual balls of wool. 6-8 small “balls” can be distinguished in Fig. 5A,C on individual focal planes (see also video 4 with the compiled animations of image parts 5A-D). Additional examples are shown in Fig. 6 (and video 5 with compiled animations of image parts 6A,C,D). The enhanced fibrin labeling observed around these platelets, compared to those in a typical constrained clot (Fig. 2), may be due to the compaction of free, unconstrained fibrin fibers present in the 2D assay. This may suggest that fibers are compacted in multiple loops around platelet bulbs during the progression of their formation. However, the current imaging resolution is insufficient to fully resolve the architecture of these densely compacted fibrin structures.

**Figure 5:**
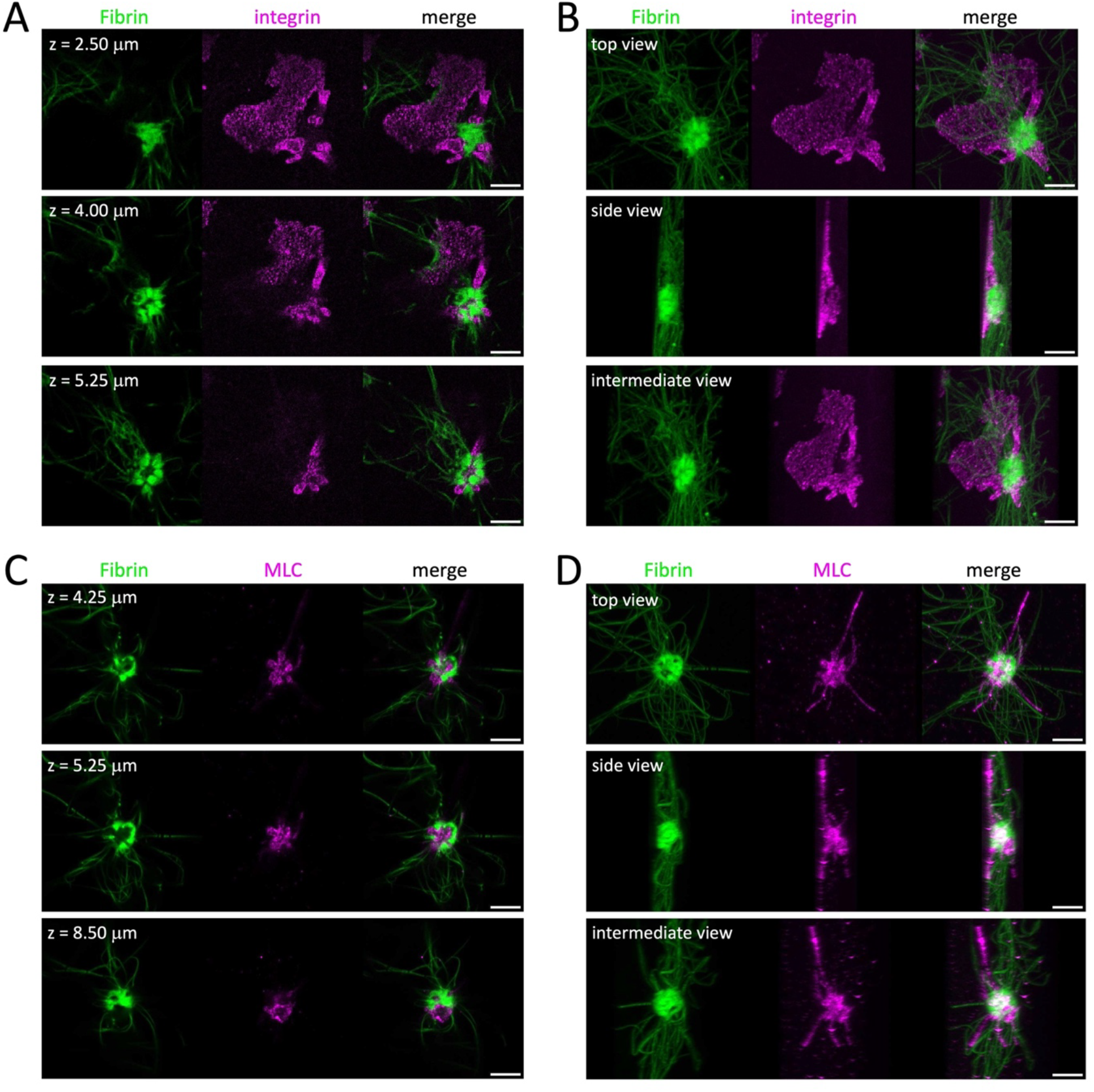
Strongly compacted fibrin fibers around the center of platelets in the 2D fiber-retraction assay. Two representative examples of platelets are shown with bulbs encircled by fibrin fibers in the 2D fiber-retraction assay (4 μl plasma/ml, see methods section) behaving similar to platelets in a clot (see also video 4 with compiled animations of image parts A-D). Samples were stained for the αIIb integrin subunit (A, B) or the myosin light chain (MLC; C,D) and processed for expansion (scale bars 10 μm = 2.5 μm after correction for expansion, indicated z-levels are not corrected for expansion). Experiment was repeated 4x with blood from different donors with a total of 18 image stacks acquired for fibrin/integrin and 62 for fibrin/myosin staining. A)Three different focal planes from an image stack showing two platelets, one spread and one with bulbs and attached, compacted fibrin fibers. B)3D image reconstruction of the image stack shown in A (34 planes, step size 0.25 μm), represented are three different view angles. C)Three different focal planes from an image stack illustrating another platelet with bulbs and attached, rolled-up, compacted fibrin fibers. D)3D image reconstruction of the image stack shown in C (55 planes, step size 0.25 μm), represented are three different view angles.

**Figure 6:**
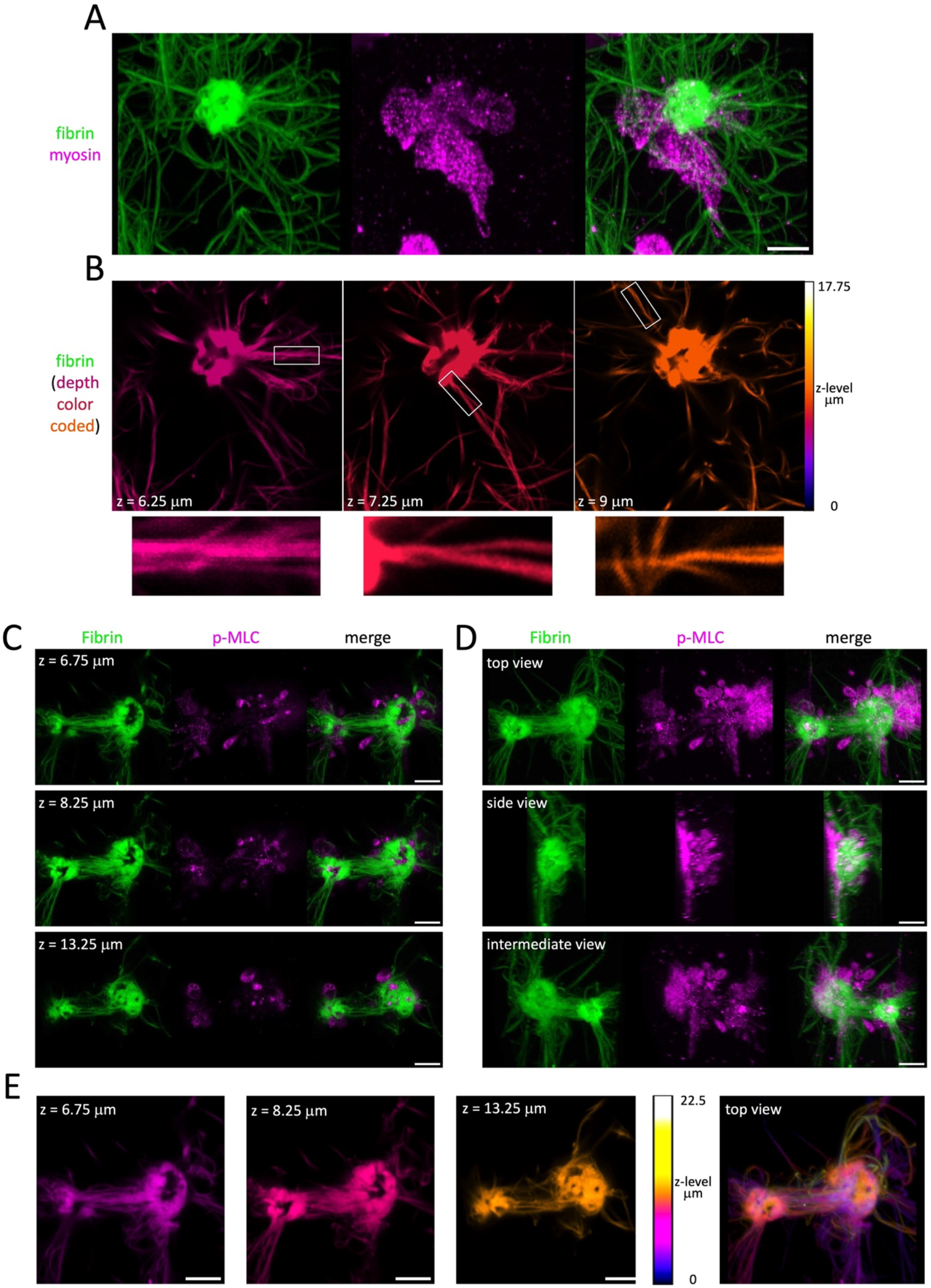
Fibrin fibers are twisted in the proximity of platelets. Two representative examples of platelets with compacted fibrin fibers are shown in the 2D fiber-retraction assay (4 μl plasma/ml, see also video 5 with compiled animations of image parts A,C,D). Samples were stained for myosin II (A, B) or phosphorylated myosin light chain (p-MLC; C-E) and processed for expansion (scale bars 10 μm = 2.5 μm after correction for expansion, indicated z-levels are not corrected for expansion). Please note also the strong fiber accumulation around the platelets similar to platelets shown in figure 5. A)3D image reconstruction of an image stack (72 planes, step size 0.25 μm) illustrating a platelet with bulbs and wound-up compacted fibrin fibers. B)Three different focal planes (z-level as indicated) from the image stack shown in A, depth color-coded. Rectangles indicate twisted fibrin fibers and a corresponding zoom is shown below. C) Three different focal planes (z-level as indicated) from an image stack illustrating two platelets with bulbs and attached, wound-up, compacted fibrin fibers. Twisted thick fiber bundles are observed between the two platelets. D) 3D image reconstruction of the image stack (91 planes, step size 0.25 μm) shown in C, represented are three different view angles. E) The same focal planes as in C and the top view of the 3D image reconstruction shown in D of the fibrin fibers, depth color-coded to better distinguish the twist of the thick fiber bundles between the two platelets.

We observe local twisting of fibers in the vicinity of platelets. Particularly clear examples are shown in Figure 6B. These observations may suggest that formation of multiple fiber loops around platelet bulbs induces twisting of the fibers, which in turn may lead to their bundling. Fiber twists can be distinguished in most of the presented images and fiber bundles between two platelets can also be twisted (Fig. 6C-E and video 5 with compiled animations of image parts 6A,C,D).

### A ring-like organization of fibrin fiber patches can be observed at the center of spread platelets

We next increased the plasma volume (7μl/ml instead of 4μl/ml) in order to enhance the amount of formed fibrin fibers. This modification led to the formation of a fragile fibrin gel, which was partially lost during the fixation or staining steps. As a result, only very few fibers remained attached to the platelets or to the coverslip.

The residual fibrin fluorescence revealed a ring-like arrangement of fibrin patches at the platelet center. This fibrin “rosette” co-localized with a circular actin organization, forming a gearwheel-like pattern with a compacted fibrin mass at the inner edge of the circle. The actin cytoskeleton was either composed of radial and transverse fibers (Fig. 7 A,B and video 6 with compiled animations of image parts 7A-D) or radial fibers and actin nodules (Fig. 7C and video 6) depending on the degree of platelet spreading.^14,15^ A faint rosette with a compact fibrin mass and an attached fibrin fiber is shown in figure 7D and video 6.

**Figure 7:**
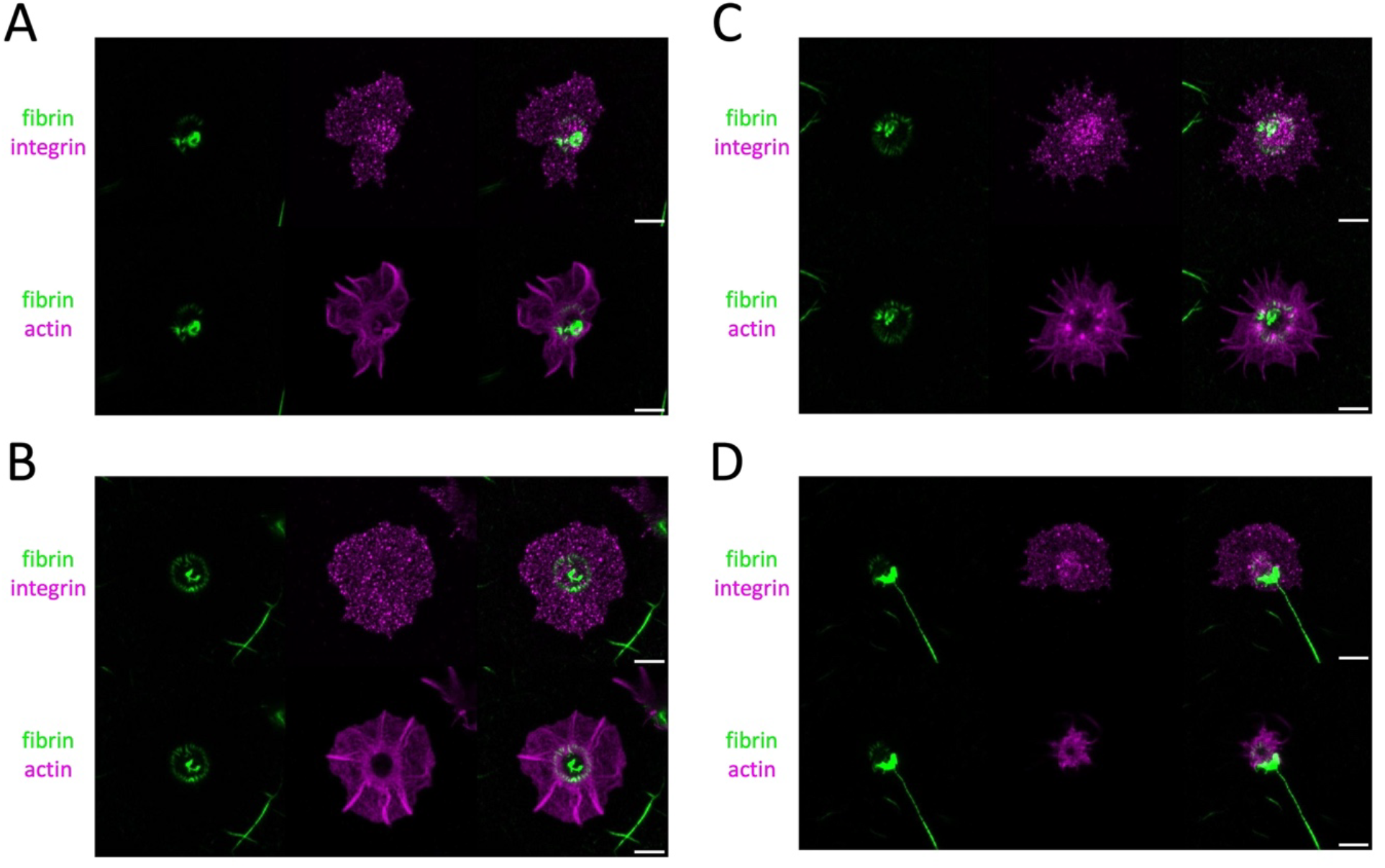
A fibrin rosette is located close to a ring-like actin organization in spread platelets. Four representative examples (out of n=20 acquisitions, experiment repeated 4 times using blood from different donors) of a fibrin rosette associated with spread platelets in the 2D fiber-retraction assay (7 μl plasma per ml PBS, see methods section) are shown (see also video 6 with compiled animations of image parts A-D). Samples were stained for the αIIb integrin subunit as well as for actin and processed for expansion (scale bars 10 μm = 2.5 μm after correction for expansion). A)3D image reconstruction of an image stack (30 planes, step size 0.33 μm) of fibrin fibers (green) and integrin or actin staining (upper and lower panels respectively, magenta). B)Similar example as in A (17 planes, step size 0.33 μm). C) Another example of a fibrin rosette with intercalated actin nodules (34 planes, step size 0.33 μm). D) An example showing a long fibrin fiber associated with the fibrin rosette (35 planes, step size 0.38 μm).

We then investigated the localization of myosin II under these conditions and found that it is distributed throughout the platelet. However, a denser ring-like organization of myosin is present in the center of the spread platelet, closely associated with the fibrin rosette (Fig. 8 and video 7 with compiled animations of image parts 8A-F). Some myosin spots are intercalated with compacted fibrin (Fig.8 A, focal plane at z=7.8 μm above the culture surface), while coiled fibrin fibers are visible at the periphery and above the myosin ring structure (Fig. 8A-F). The intensity of fibrin rosettes is highly variable between platelets of different donors. Intense rosettes, as shown in figure 7, were seen for platelets from two donors (approximately 3-5% of spread platelets with rosettes), while faint rosettes (Fig. 8AB) were observed for 50% of spread platelets using blood from three other donors.

**Figure 8:**
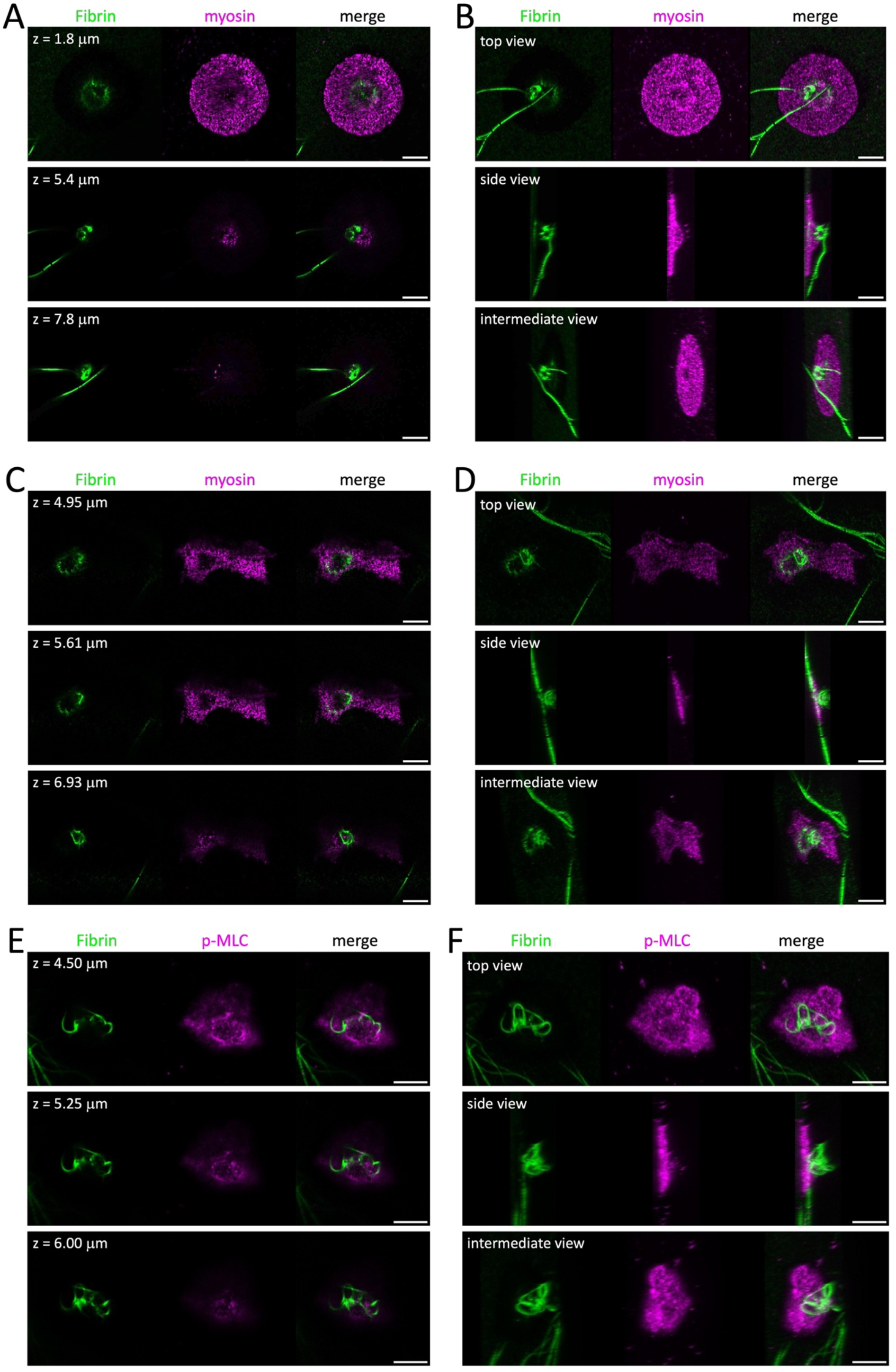
The fibrin rosette and attached fibers are in close proximity to myosin II in spread platelets. Three representative examples (out of n=62 acquisitions, experiment repeated 4 times using blood from different donors) of fibrin and myosin localization in spread platelets (see also video 7 with compiled animations of image parts A-F) in the 2D fiber-retraction assay (4 μl plasma/ml for A, B, E, F or 7 μl plasma/ml C, D; see methods section). Samples were stained for myosin (A, B, C, D) or phosphorylated myosin light chain (p-MLC; E, F) and expanded (scale bars 10 μm = 2.5 μm after correction for expansion, indicated z-levels are not corrected for expansion). A)Three different focal planes (z-levels as indicated) from an image stack showing a spread platelet and an attached fibrin fiber in the center of a faint fibrin rosette. B) 3D image reconstruction of the image stack shown in A (36 planes, step size 0.3 μm), represented are three different view angles. C) Three different focal planes from an image stack illustrating a spread platelet with a fibrin rosette and an attached, rolled-up fibrin fiber. D) 3D image reconstruction of the image stack shown in C (31 planes, step size 0.33 μm), represented are three different view angles. E) Three different focal planes from an image stack illustrating a spread platelet with attached, rolled-up fibrin fibers. F) 3D image reconstruction of the image stack shown in E (54 planes, step size 0.25 μm), represented are three different view angles.

### Modeling platelet-mediated winding-up of fibrin fibers

To address the mechanical aspects of the observed platelet-mediated fibrin fiber organizations, we developed a model based on our experimental observations, which suggests that platelets may be capable of winding fibrin fibers around their bulbs. We first quantified the number of platelet extensions. Each platelet in a clot showed 7-10 bulbs/filopodia (Fig. 9A, C). Similarly, platelets in the 2D fiber-retraction assay, which have formed bulbs and filopodia and show an accumulation of fibers around the base of their bulbs have also 7-10 bulbs/filopodia (Fig. 9B, C). Fiber curvature was determined using transmission electron microscopy (TEM) and fluorescent (FLUO) images. Curvatures were similar for both image types (mean values for κ_TEM_=10.09 ± 6.6 μm^-1^ and κ_FLUO_=6.77 ± 4.7μm^-1^) with differences likely due to the higher resolution of TEM (Fig. 9D-F).

**Figure 9:**
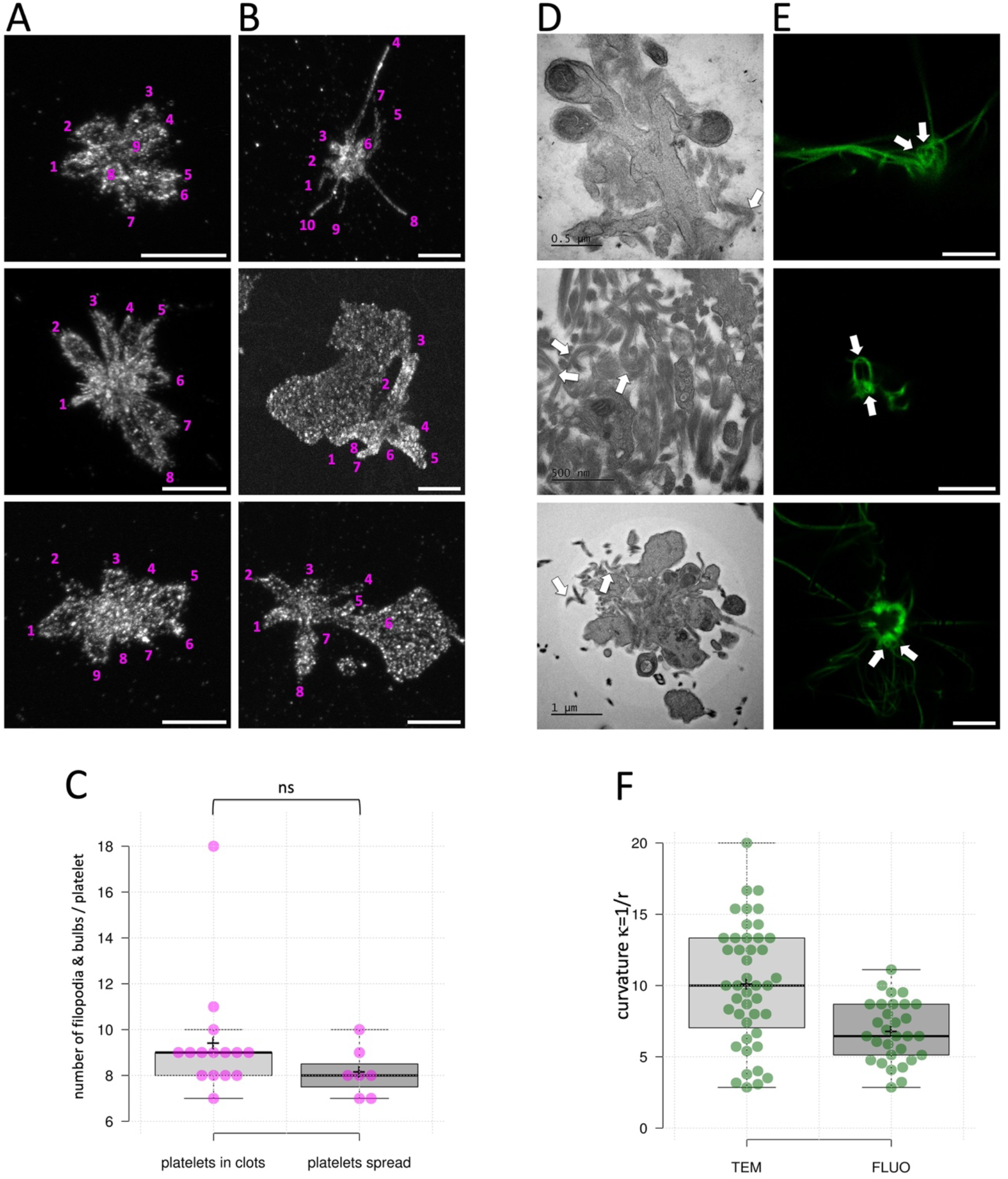
Determination of platelet protrusions and fibrin fiber curvatures in proximity of platelets. A)Three examples of platelets within a retracted clot stained for the aIIb integrin subunit (including platelets shown in figure 2). Maximum intensity projections (MIPs) are shown, the number of platelet extensions is indicated and has been determined manually by scrolling through the image z-stack. Scale bars 10 μm = 2.5 μm after correction for expansion. B)Three examples of platelets after 2D-fiber retraction (including platelets shown in figure 5) and stained for the MLC (upper panel) or the aIIb integrin subunit (middle and lower panel). MIPs are shown, the number of platelet extensions is indicated and has been determined manually by scrolling through the image z-stack. Scale bars 10 μm = 2.5 μm after correction for expansion. C)The comparison of all data points for platelets after 3D clot retraction or after 2D fiber-retraction does not show a statistically significant difference between protrusions of platelets in a clot or attached to a glass surface (Mann-Whitney U test). D-F: The degree of fiber curvature in examples shown in D and E is determined as κ=1/r using the radius of a circle fitting the region of strongest curvature of the fibers. D)Three examples of TEM images used to determine the curvature of fibrin fibers close to platelets within a clot (measured examples are indicated by white arrows). E)Three examples of fluorescent images of fibers wound around platelets attached to a glass surface (including platelets shown in figure 5). A single focal plane of each image stack is shown (scale bars 10 μm = 2.5 μm after correction for expansion; measured examples are indicated by white arrows). F) Comparison of all data points obtained using TEM images and fluorescent images (curvature in κ μm^-1^). For quantification, 31 EM acquisitions of three clots from different donors were used as well as 12 Fluo acquisitions of platelets from 4 different donors.

Our model incorporates these observations, published parameters (Table 1, methods section) and builds on the framework established by Bershadsky and colleagues.^16^ Bershadsky’s team demonstrated that in fibroblasts, cultured on circular substrates, the actin cytoskeleton is composed of radial actin fibers extending towards the cell center and circular transverse fibers enriched in myosin-IIA. Individual platelets spread on a glass surface have a similar phenotype and cytoskeletal organization. Live-cell imaging in the study by Bershadsky et al. revealed that the transverse fibers slide centripetally along the radial fibers in a myosin dependent process, generating a chiral swirling motion directed towards the cell center. They also show that fibronectin-coated beads bound to integrins on the cell surface are dragged along with the intracellular swirling of the cytoskeleton. Thus, fibrin fibers bound to integrins at the platelet plasma membrane may experience a similar rotational motion around the pseudo-nucleus. Chiral rotations of the cytoskeleton are not only observed at the level of entire cells, but also in subcellular regions, such as individual filopodia of the neuronal growth cone.^17,18^ Similarly, chiral rotations of the cytoskeleton may be present in individual bulbs of platelets in a clot.

**Table 1.** The model parameters used in the simulation. CS – current study.

| Parameter | Value | Citation |
| --- | --- | --- |
| $R$, the bulb radius | 400 nm | CS |
| Bulb length | 1500 nm | CS |
| $H$, the height of the binding sites on integrins for fiber nodes from the bulb surface | 20 nm | 42 |
| $L$, initial distance between modeled nodes in fibrin fiber | 2×45 nm<br>(2 fibrin molecules) | 43 |
| $\alpha$, angle between fiber and X-axis in 3D coordinates | $\pi/4$ or $\pi/2.5$ | CS |
| Number of nodes | 60 | CS |
| $D$, distance from the top of the bulb to the point where the node 0 was bound at $T = 0$ | 500 nm | CS |
| $\Delta t$, time step | 0.00015 $\mu s$ | CS |
| $Rate$, rate of downstream movement of integrins | 1 nm/$\mu s$ | CS |
| Rate of swirling | $\pi/500$ rad/$\mu s$ | CS |
| Temperature, $T$ | 300 K | CS |
| Drag coefficient for the node, $\eta$ | $3.77 \times 10^{-3}$ nN*$\mu s/nm$ | 24 |
| Stiffness of fibrin fiber, $K_r$ | 11.5 nN/nm | 25 |

Along this line, we aimed to establish a model which could reproduce the experimentally observed fibrin organization around the base of platelet bulbs due to the hypothetical swirling dynamics of the cytoskeleton and the associated receptors. Our simulations are based on fibrin fibers anchored to integrins in the plasma membrane of platelet bulbs and show that in principle cytoskeletal motions within each bulb can drive integrins, and thus the attached fibrin fibers, towards the platelet center, wrapping the fibers around the bulbs (Fig. 10A and video 8). The efficiency of loop formation depends on the initial fiber position (Fig. 10B-D). The process effectively winds long fibers into compact loops with a radius of ~420-nm around the bulbs (Fig. 10D) and the loop radius is determined by the radius of the bulb (equal to 400 nm in the model).

**Figure 10:**
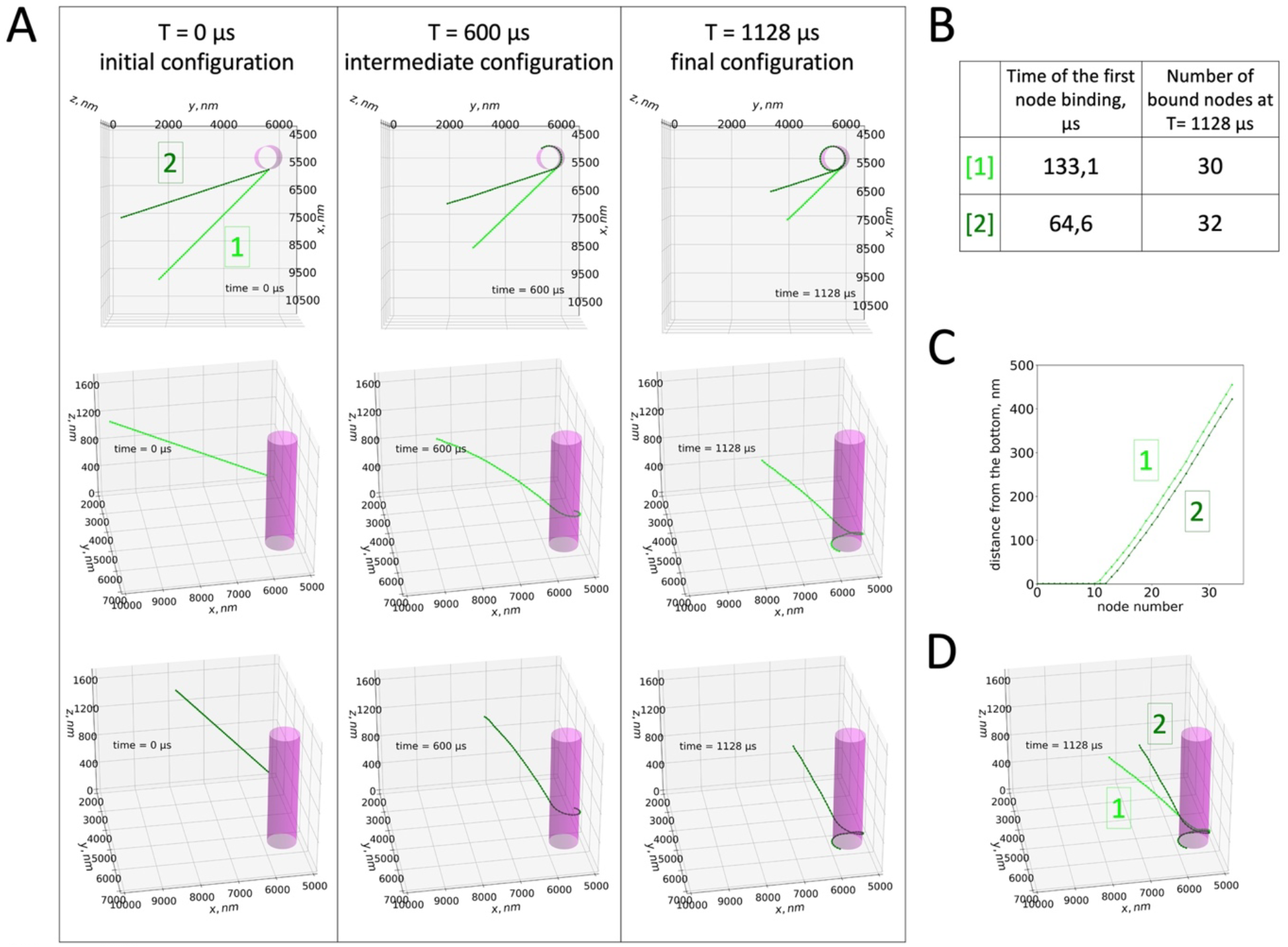
Simulation of platelet mediated winding-up of fibrin fibers. A) Time evolution of the fibrin fiber configuration around the platelet bulb. Two different angles between the fiber and the x-axis were used ([1]: α = π/4, lime color; [2]: α = π/2.5, dark green color). Left: initial positions, top view and side view. Center: intermediate position, top view and side view. The fibrin fiber began to form a loop around the bulb. Right: final position, top view and side view. A compact fibrin loop was formed around the bottom of the bulb (see also video 8). B) Time when the first node bound to the bulb and thus the formation of the fibrin loop started. The fibrin winding-up started earlier for a larger angle between fiber and x-axis (and consequently smaller angle between fiber and tangent line at the point of the node 0 binding). The number of nodes that were bound to the bulb at the end of the simulation was slightly higher for larger α. Thus, larger α led to a more compact fibrin fiber organization. C)The distance between nodes and the bottom of bulb along z-axis. The distance was smaller for larger α and the fibrin organization was slightly more compact. D)Comparison of final fiber configurations around the bulb for two different α.

In one set of calculations, we switched off the cytoskeleton swirling motion (from Table 1, rate of swirling = 0). In this case, no fibrin loop around the bulb base was formed (supplementary Fig. 1). Thus, it might be concluded that in the proposed system the cytoskeleton swirling is necessary for the fibrin loop formation.

### Transmission electron microscopy (TEM) provides evidence of curved and cross-sectioned fibrin fibers at the base of platelet bulbs

To confront the results obtained by expansion microscopy and model simulations with those using TEM, we analyzed TEM images of platelets and fibrin fibers within a clot. We observed numerous instances of cross-sectioned fibrin fibers with a distance between 200-800 nm (similar range as our model predictions) at the base of bulbs as illustrated in Figure 11. This figure also shows two examples suggesting fibers that curve along bulbs from top to bottom and fibers running along a long, thin filopodium.

**Figure 11:**
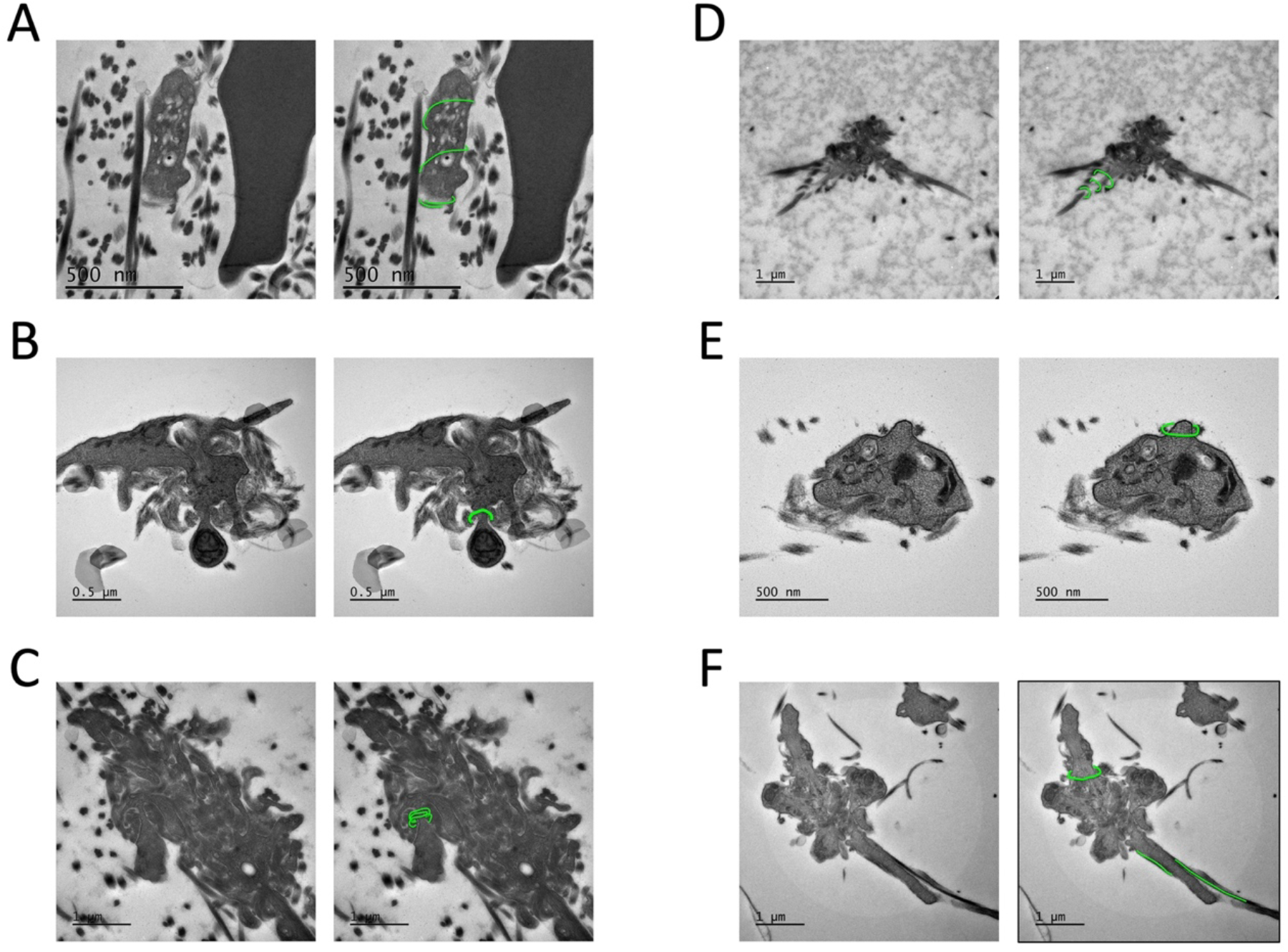
Features predicted by the model/simulation are observed on TEM images. Interpreted, potential fiber organizations are highlighted in green on an identical image on the right. A)Fibrin fibers appear to encircle a bulb. B)Cross sectioned fibrin fibers at the base of a bulb, which potentially curve around the bulb. C)Several cross sectioned fibers at the base of a bulb, which may surround the bulb. D)Several cross sectioned fibers along a platelet extension potentially forming a spiral. E)Cross sectioned fibrin fibers around a forming bulb. F)Cross sectioned fibers at the base of a bulb and two fibers along a filopodial extension.

### Live imaging of platelet-mediated reorganization and compaction of fibrin fibers

We performed live-cell video microscopy using platelets from three different donors to observe how platelets orchestrate fibrin fiber organizations (Fig. 12 and video 9 with compiled timelapse acquisitions associated to image parts 12A,B,C,E). In many cases, fibrin fibers were already accumulated above platelets at the start of imaging, with minimal further fiber movements or compactions; likely due to fibers binding to each other, to the dish or to the platelet surface (Fig. 12A). However, dynamic fiber movements, both clockwise and counterclockwise, were observed above two platelets present within the rectangle indicated in figure 12A (zoom images lower panel). An arrow in Fig. 12A marks the formation of a kink in a fibrin fiber. For the second experiment (blood from a different donor), we reduced the incubation temperature to 21°C in order to slow down the platelet and fiber movements. In the rectangle indicated in figure 12B, a fibrin fiber gets progressively curved and eventually ruptures (at time point 00:48). The resulting free end of the fiber then undergoes a clockwise rotation (viewed from below, zoom images lower panel). Two other platelets producing a fiber densification are marked by an arrow in figure 12B. One of these platelets shows a counterclockwise rotation (viewed from below) of the central part of the platelet during the fiber densification process. A third experiment (platelets from a third donor), again performed at room temperature, shows the rotational (counterclockwise, viewed from below) movement of the pseudo-nucleus and alongside the winding-up of fibers around this bulbous protrusion. This apparently generates strong tension that pulls the pseudo-nucleus towards the edge of the spread platelet, where the movement ceases (Fig. 12C, zoom images lower panel). These fibrin fiber movements occurred without visible filopodia-fiber interactions, suggesting that the bound fibers are dragged by intracellular movements of the cytoskeleton. Since still images cannot fully illustrate the fiber movements or the rotation of the platelet center, we invite the reader to consult the accompanying time-lapse videos (video 9 with the compilation of videos associated to Fig. 12A-C,E). After the time-lapse acquisitions of figures 12B and C, samples were fixed and stained for the αIIb integrin subunit to localize the position of platelets on the dish surface (Fig. 12D). The final fibrin fiber organization around platelets in these live videos of the 2D fiber-retraction assays closely resembles the arrangement of fibers around platelets in a constrained clot (Fig. 1C), except that some platelets are completely spread on the surface with no or very few fiber contacts. The platelets, which had formed bulbs and were partially spread on the dish surface, were surrounded by densely packed fiber accumulations reminiscent of the fibrin fiber network with fibrin nodes colocalizing at platelet positions (Fig. 1C).

**Figure 12:**
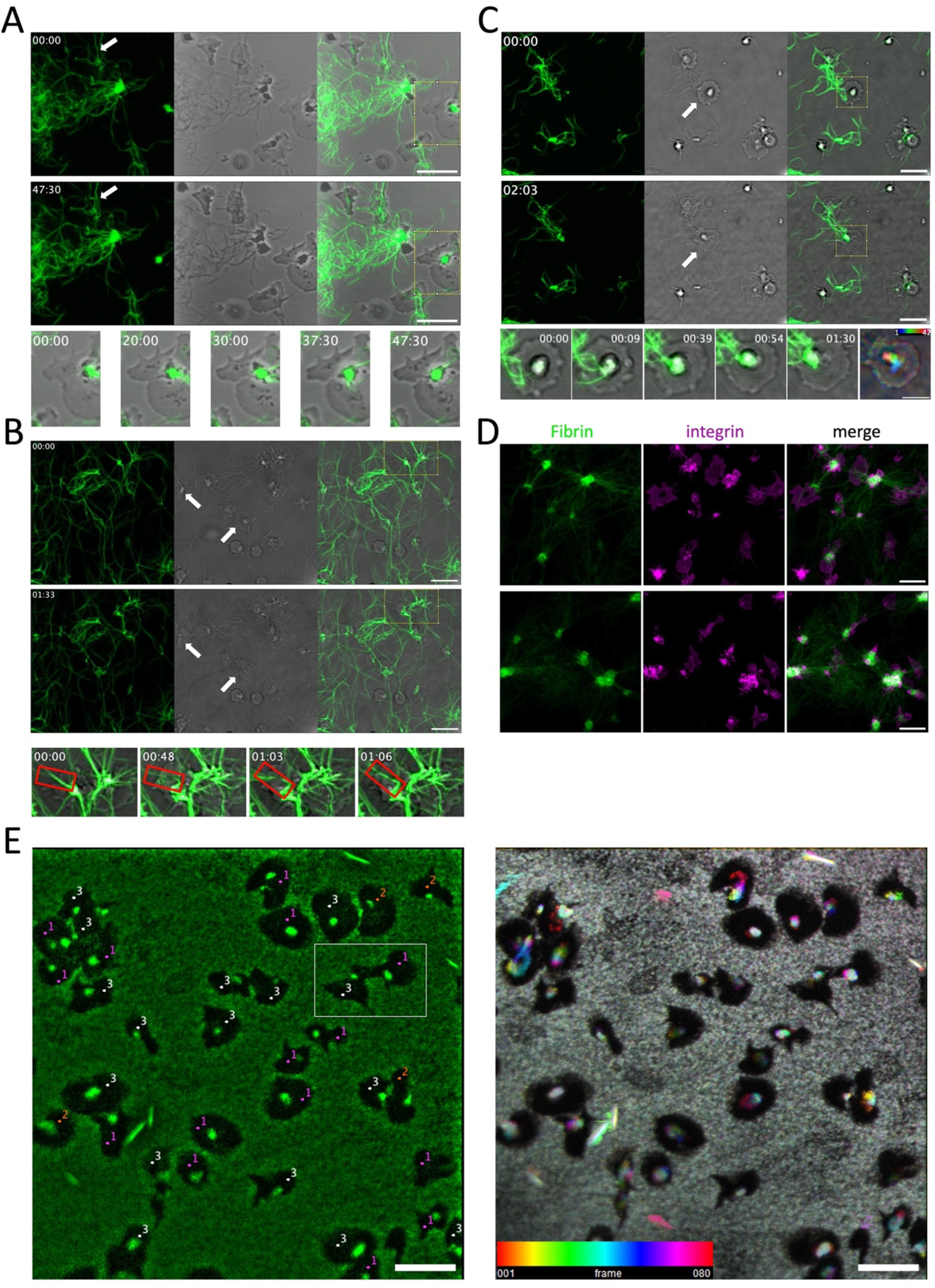
Live imaging of spread platelets winding-up fluorescent fibrin fibers. A-C: Shown are three experiments using the 2D fiber-retraction assay with blood from different donors. For each experiment, projections of four focal planes are shown corresponding to the fluorescent fibrin fibers (green), transmission and merged images captured at the first and last time points of the timelapse videos. A)Washed platelets were adjusted to 5×10^6^ platelets per 2 ml PBS and 5 μl plasma was added as well as fibrinogen-Alexa 488. Thrombin was added to induce fibrin fiber formation and platelet activation. After 10 min, the suspension was transferred into a petri dish (WPI Fluoro), centrifuged and installed in the microscope incubator at 37°C to start image acquisition (video 9, upper panel). An arrow in the fluorescent images indicates a forming kink in a fibrin fiber (scale bar 10 μm). The rectangle in the merged images indicates a region where rotational fiber movements take place. Zoom images of this region at different time points are shown below. B)The second experiment was performed under conditions as described in A except that the petri dish was kept at room temperature to slow down the platelet mediated fiber reorganizations (scale bar 10 μm). Arrows in the transmission images indicate two platelets that compact fibrin fibers. The rectangle in the merged images indicates a region where a fibrin fiber is coiled by the platelet and finally ruptures causing the broken fiber to rotate (video 9, second panel). Zoom images of this region at different time points are shown below. C)Platelets of a third donor (not washed, i.e., 5 μl PRP with 4×10^6^ platelets) were added to 2 ml PBS as well as fibrinogen-Alexa 488 and the experiment was continued as in B (scale bar 10 μm). The square in the merged images indicates a region where fibrin fibers are getting coiled and compacted around the pseudo-nucleus of the spread platelet (video 9, third panel). Zoom images of this region at different time points are shown below. The last zoomed image shows a temporal color-coded projection of the transmission channel to illustrate the rotational movement of the pseudo-nucleus (scale bar 5 μm). D)Samples shown in B and C were fixed 3h after the start of time-lapse acquisitions and stained for the aIIb integrin subunit. Shown are fibrin fibers (green) integrin staining (magenta) and the merge (samples B and C, upper and lower panels, respectively; scale bar 10 μm). E) Preformed fluorescent fibrin fibers were prepared in the absence of platelets. PRP was adjusted to 2.5×10^6^ platelets per ml with PBS and 1800 μl was transferred into a petri dish (WPI Fluoro). The dish was centrifuged to allow spreading of the platelets and the supernatant was replaced by the preformed fibrin fibers. The dish was installed in the microscope incubator to start image acquisition (video 9, last panel). The experiment was performed twice with blood of the same donor. The last fluorescent image of the time-lapse video is shown (left image). Fifteen of the fibrin accumulations in the middle of the platelets are rotating in a counterclockwise direction ( indicated as “1”), four rotate clockwise (“2”) and for 16 no clearly visible turn is observed (“3”). A projection of all fluorescent time points, temporal color-coded using the LUT spectral, is shown. White corresponds to the sum of all the colors at each time point, meaning no movement during the time-lapse video (right image; scale bars 10 μm).

To minimize pre-imaging fiber buildup, we next added preformed fibrin fibers to pre-spread platelets and started imaging immediately. Under these conditions, the glass surface is coated and only very few fibers are present. However, a central fibrin accumulation is consistently observed on each platelet, potentially serving as an initiation site for subsequent fibrin fiber organization. Among 35 platelets analyzed, 15 displayed a central fibrin mass rotating counterclockwise (viewed from below), 4 rotating clockwise, and 16 showed no consistent directional movement (Fig. 12E). In some instances, the rotating complexes stalled and stopped moving. These observations support the idea that cytoskeletal swirling could drive fibrin reorganization above spread platelets.

The fiber compaction process mediated by three platelets (two in figure 12B, one in figure 12C, indicated by arrows in the transmission images) was quantified and a fibrin accumulation of a fiber length equivalent to 20, 30 and 70 μm (accumulation rate 1, 0.51 and 1.61 μm/min, respectively), was estimated. Accumulation ceased at an approximate total fibrin fiber length of 80-90 μm (Fig. 13 A-C).

**Figure 13:**
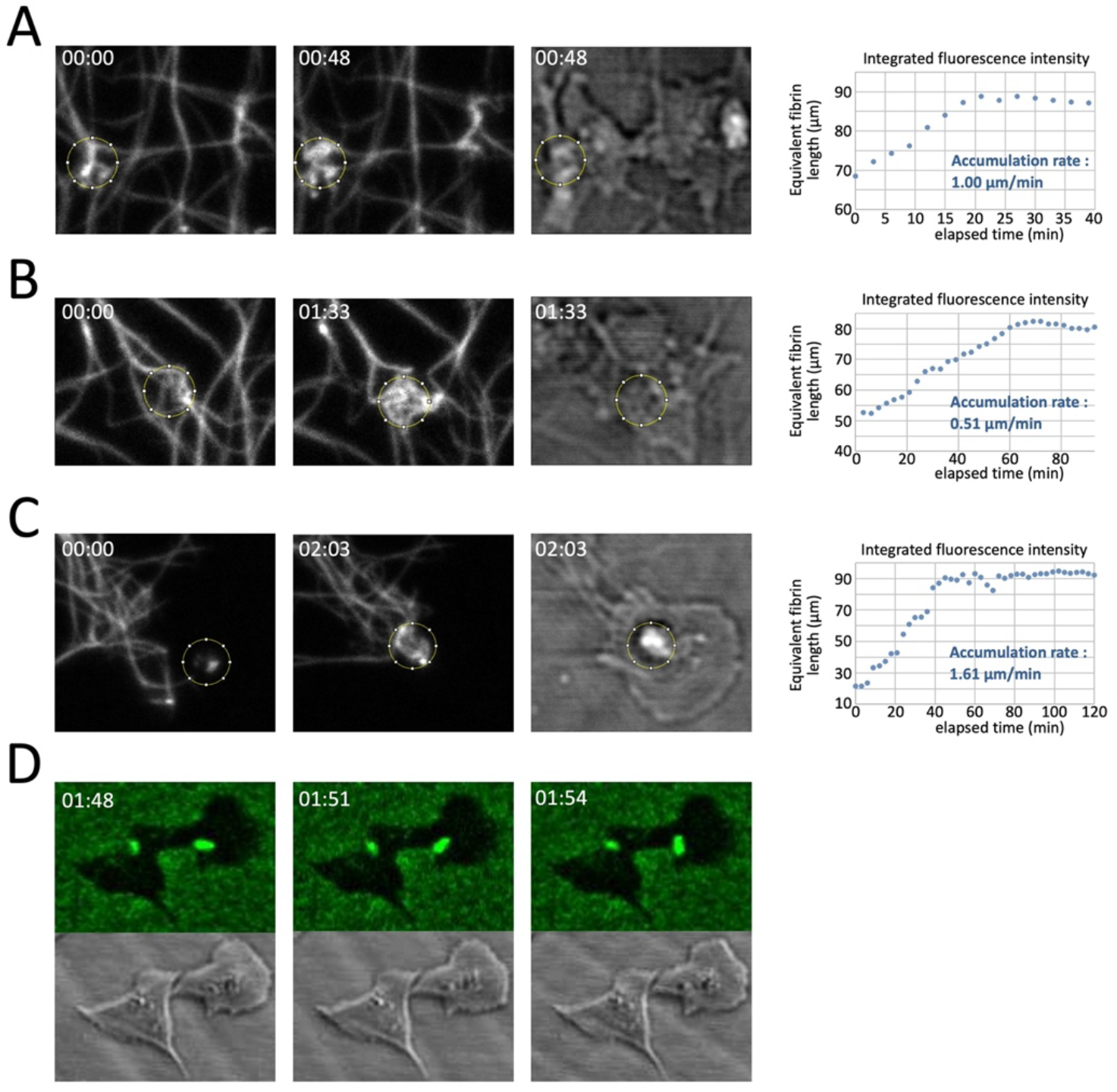
Quantification of platelet induced fiber compaction and angular rotation using the time lapse videos shown in figure 12 (video 9) A-C: Quantification of fibrin accumulations induced by 3 platelets during time-lapse confocal 3D video microscopy. In each row first and last frames of the series are shown as Z-projections of fibrin fluorescence, followed by the transmitted light image of the platelet in the last frame. The integrated fluorescence intensity in the ROI was quantified as the sum of all Z-planes, and normalized to the intensity of a 1 μm long segment of a single fibrin fiber. The equivalent fibrin fiber length accumulation is assessed as a slope of the linear regression of the initial raising part. The ROI diameter is 3 μm. A)Platelet shown in figure 12B upper left side (indicated with an arrow in the transmission image) B)Platelet shown in figure 12B middle (indicated with an arrow in the transmission image) C)Platelet shown in figure 12C (indicated with an arrow in the transmission image) D)Quantification of the angular rotation speed of a fibrin mass attached to a spread platelet during time-lapse confocal 2D video microscopy (region indicated by a rectangle in Fig.12E, left image). The right platelet presents a fibrin mass attached to the platelet surface, which undergoes a counter-clock wise rotation. Three consecutive frames of the series are shown (time interval 3 min, total duration 240 min, elapsed time as indicated), where rotation speed was maximal and the transmitted light images are shown below. The transmission images show two rotating centers present in this platelet and the fibrin mass is turning around the left center which is within the elongated spread part of the platelet indicating that lamellar protrusions may acquire a new subcellular cytoskeletal unit similar to individual bulbs of platelets in a clot. The total number of turns of this fibrin mass was about 3.3, resulting in the average angular speed of 5°/min with short-term bursts up to 18°/min.

The angular rotation speed of a fibrin mass around a cellular turning center was calculated to be 5°/min with short-term bursts of 18°/min (right platelet within the rectangle indicated in figure 12E, left image). The transmission images show two rotating centers present in this platelet and the fibrin mass is turning around the left center which is within the elongated spread part of the platelet. It is therefore possible that in lamellar protrusions a subcellular cytoskeletal unit is formed, which undergoes individual rotational movements. Formation of transient secondary centers can be seen in other platelets of this video and in a study by Gaertner et al. showing a video of platelets migrating on a fibrinogen substrate which they collect during the migration process (see video mmc6 of their publication^19^).

## Discussion

Using a 2D fiber-retraction assay, we identified a previously unrecognized capacity of platelets to compact fibrin fibers into a small volume possibly by winding them into coiled structures during platelet induced fiber retractions. To our knowledge the only example of natural fiber compaction is coiling of DNA molecules.^20^ In contrast to intracellular, enzyme-mediated DNA compaction, extracellular fibrin fiber compaction requires the cellular activity of platelets. Our investigation began with the observation that platelets become labeled with fluorescent fibrinogen during clot retraction as previously shown by Brzoska et al.^21^ They demonstrated that fibrin binding to thrombin activated platelets depends on αIIbβ3 integrins and the actin cytoskeleton. While fibrin fiber retraction by platelets in a clot has been attributed to non-muscle myosin II,^22,23^ the mechanism underlying fiber organization and compaction has remained unclear.

One key mechanism, revealed in previous studies, is the extension of platelet filopodia that bind to fibrin fibers and pull them inwards in a hand-over-hand motion.^10^ This is clearly the main process by which platelets retract fibrin fibers in a clot. However, this mechanism alone cannot explain the cage-like fiber organization observed around platelets in a constrained clot (Fig. 2). Our findings introduce an additional, distinct mechanism by which platelets can organize fibrin fibers. Platelets compact fibrin fibers into dense, ball-like structures, which is an efficient form of compaction (Fig. 4-6). Thus, our results reveal an additional mechanistic aspect of platelet mediated fiber retraction beyond those described by Michael et al.^24^ and Kliuchnikov et al.,^25^ so far observed only in the 2D fiber-retraction assay. Nevertheless, the platelet morphology and the organization of fibers around them after the retraction process closely resemble those of platelets in a clot and both, 2D and 3D fiber retractions, depend on myosin actions. Thus, it is unlikely that fibrin fibers, which present a helical twist and an inherent tension,^13,26^ could spontaneously form these ball-like structures independently of platelet activities.

The 2D fiber-retraction assay may recapitulate early post-injury events, such as activation of platelets in suspension triggered by agonists released from neighboring platelets, ultimately leading to the spreading of activated platelets onto the subendothelial matrix to anchor the developing clot at the site of vessel injury. This assay revealed two key observations:

First, when many fibers are present, some platelets spread with the typical fried-egg phenotype and coil up fibers above their pseudo-nucleus (Fig. 4). Others form large bulbs in several directions, similar to platelets in a clot, and compacted fibers are observed around the base of each bulb (Fig. 5, 6). Under these unconstrained 2D conditions, fiber accumulation around the bulbs is much higher than in the case of a constrained 3D clot (Fig. 2), where internal tensions limit fiber accumulation around the base of platelet bulbs. It is noteworthy to emphasize that under all assay conditions used (constrained or unconstrained clots, 2D fiber-retractions) only homogenously distributed platelets are investigated. Thus, the results only show how individual platelets and not platelet aggregates act on fibrin fibers.

Second, when fewer fibers are retained on the culture surface, several spread platelets display a fibrin rosette around their pseudo-nucleus (Fig. 7,8) that colocalizes with actin, myosin and integrins. A dense fibrin mass is seen at the inner periphery of this rosette, sometimes with an attached fiber. This gearwheel-like architecture may represent an early priming/nucleation step of platelet-induced winding of fibrin fibers. The actin cytoskeleton in platelets exhibits a chiral organization^27^ which, combined with the action of myosin, could be responsible for the rotational motion observed in spread platelets (Fig.12 E and video 9). Integrin-bound extracellular fibers are pulled along by this vortex inducing their coiling around the pseudo-nucleus (Fig. 12C). Fibrin rosettes may fade as coiled fibers become more prominent, suggesting a transition from an initiation phase to active coiling and compaction.

### Ideas and Speculations

How this 2D “gearwheel” translates to the 3D clot environment remains unclear. One possibility is that the pseudo-nucleus of spread platelets may correspond to a bulb around which fibers are coiled and accumulate. Platelets with bulbs within a clot have been described previously.^28-31^ We wondered whether platelet bulbs might represent plasma membrane blebs similar to membrane protrusions formed by megakaryocytes during clot retraction.^32^ However, the consistent presence of 8–9 bulbs per platelet (Fig. 9A-C) shows the three-dimensional morphology of individual platelets embedded in a fibrin clot, reminiscent of the eight corners of a cube, allowing platelets to sense the clot’s 3D environment along the main directions. Unlike blebs, these bulbs often contain organelles such as mitochondria (Fig. 9D, upper image and Fig. 11B) suggesting that each bulb may have its own energy supply and actomyosin organization. This could enable swirling within each bulb and explain the observed fibrin fiber accumulation at their base. In support of this hypothesis, we have shown that radial actin fibers extending to the platelet center are present in each platelet bulb within a constrained clot (Fig. 2E). Thus, at the onset of clot retraction, each emerging bulb may act as a discrete functional unit onto which the gearwheel assembles, enabling fibrin fibers to attach and initiating their subsequent winding and compaction (scheme, Fig. 14). These bulbs are not static structures, but are constantly moving and renewing. If fibrin fibers bind to integrins on the bulb plasma membrane without subsequent transport of these adhesion complexes towards the bulb base and recycling of integrins back to the top, both bulb remodeling and fibrin network compaction would likely be impaired. The rotational cytoskeletal motion that drags attached fibers towards the bulb base cannot be interpreted as a simple winding of individual fibers, since platelets are embedded within a highly interconnected fibrin network. Instead, rotational transport of bound fibers within this network may generate constrictive forces at the bulb base, where fibers progressively accumulate. This accumulation could promote local fiber densification and thereby contribute to compaction of the fibrin network as a whole.

**Figure 14:**
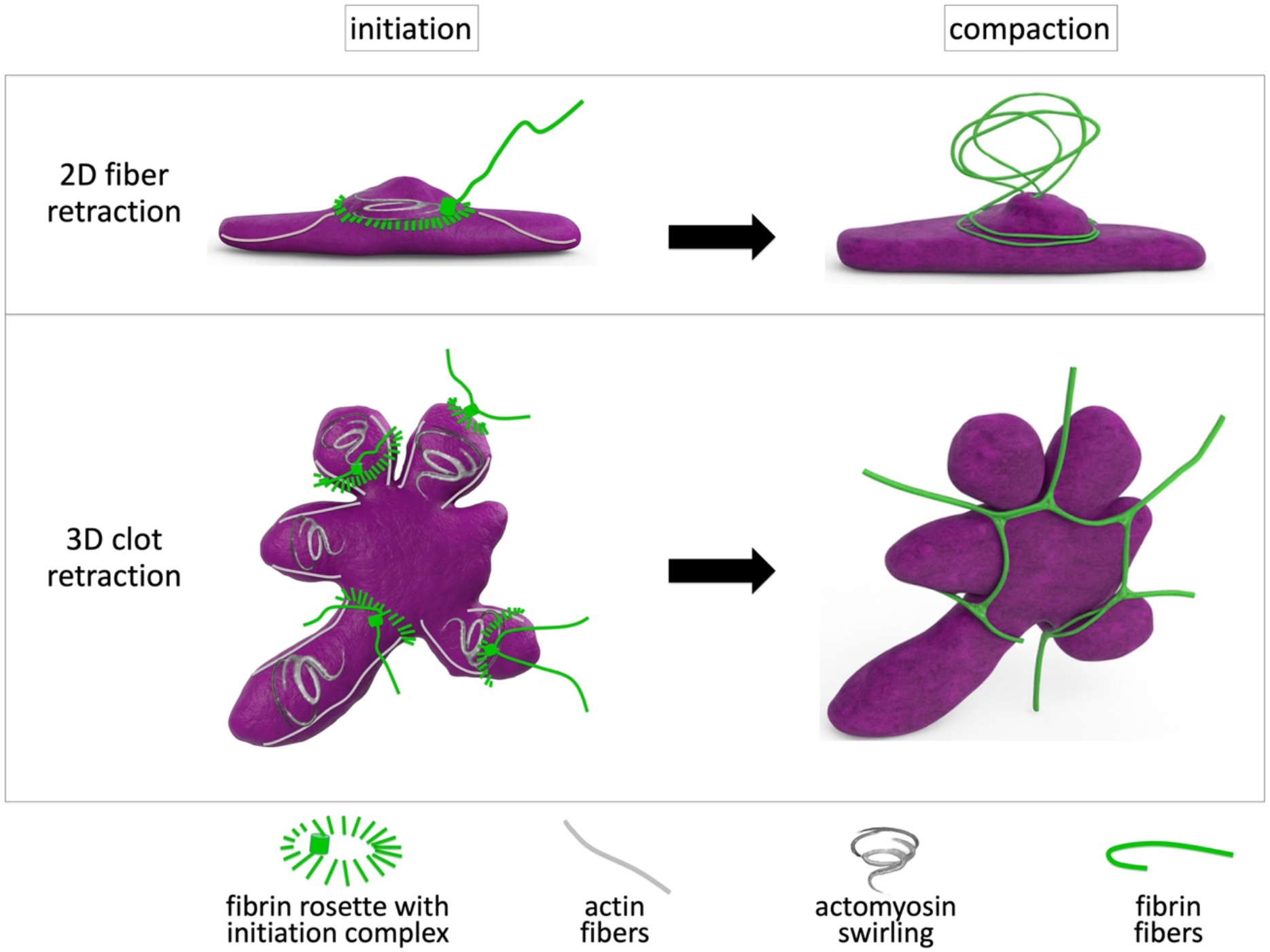
Schematic illustration of the hypothesis explaining the potential mechanism by which platelets may wind-up fibrin fibers and compact them. Represented is a platelet in the 2D fiber-retraction assay (upper part) or in a clot (lower part). At the beginning of platelet fiber interactions, a fibrin rosette is present around the pseudo-nucleus and in each bulb with a dense fiber mass at the internal periphery of the rosette possibly representing an initiation complex. Radial actin fibers extend to the center of the platelet and an actomyosin rotational movement along the radial actin fibers may lead to the winding-up and the downward sliding of the extracellular attached fibers to encircle the pseudo-nucleus or the base of the bulbs at the final state of compaction (3D rendering of the scheme was produced using an open AI).

Indeed, blebs of megakaryocytes retracting a clot may behave similarly as platelet bulbs since fibrin fibers can be seen surrounding the base of blebs forming a cage-like structure around the center of the megakaryocyte during megakaryocyte-mediated clot retraction, as observed by Kim et al. (see the supplementary video 3: blood_bld-2023-021545-mmc3 of their publication).^32^

Our live time-lapse videos of platelet-mediated fiber movements in a 2D environment support the concept of a cytoskeletal vortex that drives the coiling and compaction of extracellular attached fibers (Fig. 12A-C). Additionally, the last time-lapse video capturing the rotation of compacted fibrin clusters at the center of platelets (Fig. 12E) further suggests the presence of cytoskeletal swirling within spread platelets. During fibrin fiber formation this swirling may play a role in scanning the extracellular space for forming fibers that can be anchored to the initiation complex. The process occurs on a notably slow time scale, which could facilitate effective fiber capture by the rotating complex. However, this rate appears insufficient to account for the rapid labeling of platelets during clot formation, suggesting that regulatory mechanisms, possibly involving mechanical feedback, may be at play. Indeed, as mechanosensory cellular particles, platelets adjust their contractile forces and dynamics in response to environmental cues such as fiber tension and clot stiffness.^9,10,33,34,25^

In conclusion, our study points to a novel mechanistic aspect of platelet function: the potential active winding-up and compaction of fibrin fibers. This concept complements the existing model of platelet-mediated clot retraction^10^ and provides a new perspective on how platelets may contribute to clot architecture, mechanical stability, and wound healing.

## Materials and methods

### Reagents and antibodies

Reagents: Neutral formalin (Sigma-Aldrich, Saint-Quentin Fallavier, France), Pluronic F-127 (Sigma-Aldrich, P2443), thrombin (Sigma-Aldrich, T4648), Jasp-K4-HA (a HAK-actin probe, kind gift of Paul Guichard/Virginie Hamel),^35^ apyrase (Sigma-Aldrich, A6535), heparin (Sigma-Aldrich, H3393), PGI2 (Sigma-Aldrich, P6188), NHS-Ester DyLight 650 (Thermo Scientific) Antibodies: mouse anti-integrin αIIb (Santa-Cruz, sc-7310), monoclonal rabbit anti-HA (Cell Signaling, C29F4), rabbit anti-non muscle myosin II heavy chain (Covance, PRB-440P), rabbit anti-phospho S19 myosin light chain (Abcam, ab2480), monoclonal rabbit anti-myosin light chain 2 (Abcam, ab92721), Cy3 goat anti-mouse IgG (Jackson, 115-165-166), Cy3 goat anti-rabbit IgG (Jackson, 111-165-003), Alexa 647 goat anti-rabbit IgG (Jackson, 111-605-003)

### Preparation of human platelet rich plasma (PRP) and platelet free plasma (PFP)

Whole blood was obtained from the French blood bank (age-range 18-65, mixed genders; pooled blood from multiple donors has not been used). The IRB has authorized our institute’s research projects using human samples under the number DC-2022-4976. PRP was prepared from whole blood drawn into Na-citrate Vacutainers (3 tubes of 2.7 ml per donor). The blood is pooled into a 15 ml falcon tube and centrifuged for 12 min 400g, RT, no brake. The upper phase is the platelet rich plasma (PRP), in which platelets are distributed in a gradient (low concentration on top, high concentration at the bottom). In order to increase the platelet concentration in the final PRP, the upper 3 ml are removed and used to prepare platelet free plasma (PFP, by centrifugation at 12000g, 1 min, RT). The remaining upper phase is collected as the final PRP and the platelet concentration determined.

### Platelet washing procedure

PRP was diluted 1/4 with Tyrode/0.35% BSA buffer and platelet activation inhibitors (0.02 U/ml apyrase, 1 μm PGI2, 10 U/ml Heparin, final concentrations) were added. Samples were then centrifuged for 15 min at 900g at RT. The supernatant was aspirated, and platelets were gently resuspended in fresh Tyrode/BSA buffer, 0.02 U/ml apyrase, 1 μm PGI2. The suspension was centrifuged as before and the platelet pellet was resuspended in fresh Tyrode/BSA buffer without inhibitors.

### Clot retraction assays

We used three different clot retraction assays illustrated in supplementary figure 2.

1. Unconstrained clot retraction: See paragraph “Live video microscopy” below for method details.
2. Constrained clot retraction between two holders: In order to perform immunofluorescence staining of individual platelets within a clot, we used a clot retraction assay around two sterile inoculation loops as previously described.^36,37^ Briefly, PRP was adjusted to 1×10^8^ platelets per ml in 50% plasma/50% PBS (to reduce fibrin fiber density), fibrinogen-Alexa 488 final concentration 12.5 μg/ml and 4 μl/ml erythrocytes for color contrast were added. 400 μl of this suspension was used for clot formation by addition of thrombin (2.5U/ml final) and immediately transferred to 2 ml Eppendorf tubes coated with 2% Pluronic® and containing the inoculation loops. After different retraction times, clots were fixed for 1 h with isotonic formalin (9 volumes formalin / 1 volume 10xPBS). Clots were then washed with PBS and incubated ON at 4°C in 1 ml PBS/15% sucrose. Clots were further incubated in 1 ml PBS/15% sucrose/7.5 % gelatin at 37°C for 4 h and then inclusion blocks were formed in the same solution on ice. Blocks were then trimmed and snap frozen for 1 min in isopentane at minus 60°C and stored at minus 80°C until 14 μm thick transversal sections were cut using a cryostat.
3. Constrained clot retraction within an inoculation loop: Small clots of 30 μl platelet suspension (50% plasma/50% PBS; 1×10^7^ platelets/ml; fibrinogen-Alexa 488 final concentration 12.5 μg/ml; 4 μl/ml erythrocytes for color contrast) were induced to form within plastic inoculation loops (4 mm internal diameter, attached to a glass coverslip) by addition of thrombin to a final concentration of 2.5 U/ml as described previously.^15^ Five minutes after clot induction, clots were covered with 500 μl PBS and kept in a cell culture incubator for 10 min. Clots were then fixed with 500 μl isotonic formalin for 15 min, RT, in the dark and then kept in PBS at 4°C until further use. Forming a clot within a small inoculation loop, allowed us to do expansion microscopy without performing cryosections of larger clots, which are difficult to detach from slides during the expansion process.

### 2D fiber-retraction assay

PRP was diluted in PBS (2.5×10^6^ platelets/ml, plasma concentrations were kept at 4 or 7 μl/ml depending on the amount of fibrin fibers to be formed; see explanation below) and fibrinogen-Alexa 488 was added to a final concentration of 12.5 μg/ml. To induce fibrin fiber polymerization and platelet activation, thrombin was added (final concentration of 2.5U/ml). The suspension was gently mixed once (to prevent platelet aggregation) and after 15 min of incubation at RT in the dark, 400 μl was transferred into wells of a 24 well plate containing glass coverslips. The plate was centrifuged 3 min, 600 g, RT to allow synchronized contact of all platelets with the glass surface. The plate was then placed in a cell culture incubator for 30 min to allow platelet spreading. Samples were then fixed with isotonic formalin for 15 min at RT and then kept in PBS at 4°C until further use.

To promote fibrin fiber formation, while preventing the development of a clot, the plasma concentration must be kept low (4 μl plasma per ml PBS). This can be achieved by adding a volume of PRP containing 2.5×10^6^ platelets to 1 ml PBS, provided the platelet concentration of PRP is at least 6.25×10^8^/ml (condition used for Fig. 3, 4, 5, 6, 8ABEF). If the platelet concentration in the PRP is lower, platelets must be washed and the appropriate amount of plasma added separately.

At higher plasma concentrations (7 μl plasma per ml PBS), more fibers polymerize leading to the formation of a fragile clot which is aspirated during fixation/washing procedures. Under these conditions very few fibers are left behind. However, a fibrin rosette can be observed in the center of several spread platelets (condition used for Fig. 7, 8CD).

The 2D fiber-retraction assay followed by immunofluorescence staining and expansion has been repeated 10x with blood from different donors. A total of 18 image stacks stained for fibrin/integrin, 20 stained for fibrin/integrin/actin and 62 stained for fibrin/myosin have been acquired.

### Immunofluorescence

Fixed platelets were permeabilized with PBS/0.2% Triton X-100 for 15 min at RT and then incubated with blocking buffer (3% BSA and 10% goat serum in PBS) for 1h at RT. Coverslips were then incubated for 2 h with a primary antibody diluted in blocking buffer, then washed three times and incubated with a secondary antibody diluted in blocking buffer for 2h at RT. After three washing steps, coverslips were either mounted in Mowiol or processed for expansion.

### Expansion microscopy

Expansion microscopy was essentially performed according to Gambarotto et al.^38^ with minor modifications as described previously.^37^ Briefly, fixed, immunostained platelets or clots on coverslips (12 mm diameter) were incubated in PBS/0.7% formaldehyde/1% acrylamide over night at room temperature. The coverslip was then placed upside down onto an ice cold 35 μl drop of gelation solution (PBS/19% Na-acrylate/10% acrylamide/0.1% bisacrylamide) pipetted onto parafilm on ice immediately after addition of TEMED and ammonium persulfate both to a final concentration of 0.5%. After 5 min of incubation on ice, the samples were transferred to 37°C for 1 h. The coverslip plus gel facing up was then incubated in denaturation buffer (5.7% SDS/0.2M NaCI/50mM Tris, pH9) for 15 min at RT with gentle agitation to detach the gel from the coverslip. The gel was then transferred into an Eppendorf tube with fresh denaturation buffer and boiled for 30 min at 95°C. To expand the gel, it was placed into a large volume of H_2_0 and the water was changed twice before incubation ON at RT in fresh H_2_0 leading to 4x isometric expansion.

### Image acquisition

For image acquisition of some of the expanded samples (Fig. 2, 5CD, 6, 7, 8), a confocal microscope ConfoBright (Nikon A1R+MP) equipped with a home-made adaptive optics module was used with a 40x/1.15 NA apo long distance water immersion objective. The geometric optical aberrations were corrected both in excitation and detection light paths in the open loop mode with a deformable mirror (AlpAO DM97-15) inserted between the confocal scanning head and the microscope inverted stand (Nikon Ti2E). Local aberrations were sensed using the metrics of molecular brightness, derived from fluorescence fluctuations, and were iteratively corrected as individual Zernike modes. This configuration allowed to compensate for the refractive index mismatch, the specimen tilt and the eventual light scattering in depth of expanded samples and to bring back the confocal resolution to its diffraction limit in the vicinity of the imaged platelet.

Image acquisition of other expanded samples (Fig. 4, 5AB) was performed with a confocal microscope Dynascope (LSM 710 ConfoCor 3; Zeiss), 63×/1.4 NA Plan Apochromat objective and the acquisition software Zen 2010. The same microscope and software were used for live imaging (Fig. 12) using the Apochromat 40×/1.2 NA water-immersion objective. Axial Z-stacks (step 0.45 μm) were imaged in a time series using the avalanche photodiode detector in photon counting mode at low 488 nm excitation power (0.1% transmission) to limit photobleaching and phototoxicity, the 633 nm laser was set to 0.6% in order to obtain the transmission light images of the platelets. The pinhole was closed to 0.25 Airy units (14 μm) to increase the spatial resolution. The acquisition of transmitted light was concomitant to the fluorescence.

The FIJI software was used for image processing and analysis.^39^ No treatment that modifies the raw images was applied. FIJI software was used for the Maximum Intensity Projection (MIP) images, 3D reconstructions of image stacks and for the depth-color coded projection image using the rainbow LUT (Fig. 4D), the color code in the HSB space (Fig. 4E) as well as for the depth-color coded images fire LUT (Fig. 6) and the temporal-color coded image spectral LUT (Fig.12E).

### Transmission electron microscopy (TEM)

Clots were formed around two holders as described previously.^36^ Briefly, PRP was adjusted with PBS to 10 % plasma, 1×10^8^ platelets/ml and 4ml erythrocytes per ml were added for color contrast. Clot formation was induced by addition of thrombin (final concentration of 2.5 U/ml). After 1h of retraction at 37°C, clots were fixed using 1,5 ml of 2,5 % glutaraldehyde for 1h at RT. After fixation, a contrast-enhancing step was performed by incubating the samples in a solution containing 1.5% potassium ferrocyanide and 1% osmium tetroxide in 0.1 M sodium cacodylate buffer. Following this, the samples were embedded in EPON resin according to the protocol described by Eckly et al.^40^ Thin sections (100 nm) were cut and stained with uranyl acetate and lead citrate and examined using a Jeol 1200 Plus transmission electron microscope operating at 120 kV.

### Live video microscopy

- Unconstrained clot retraction (Fig. 1B): Thrombin (final concentration of 2.5 U/ml) was added to 0.4 ml of a platelet suspension (50% plasma/50% PBS; 1×10^8^ platelets/ml; fibrinogen-Alexa 488 final concentration 12.5 μg/ml; 4 μl/ml erythrocytes for color contrast) to induce clot formation. After quick and gentle mixing, the suspension was transferred immediately into a well (8-well Lab-Tek, coated 2% Pluronic® to prevent clot attachment to well walls). Acquisition was started immediately at 100 μm above the bottom of the well using the confocal microscope ConfoBright (for microscope details see paragraph image acquisition).
- 2D fiber-retraction assay (Fig. 12A-D): Platelets (washed Fig. 12AB or not Fig. 12C) were adjusted to 2.5×10^6^ platelets/ml PBS and 2.5 μl plasma was added and fibrinogen-Alexa488 (final concentration of 12.5 μg/ml). 15 minutes after thrombin induced fibrin fiber formation and platelet activation, 1800 μl of the suspension was transferred into a petri dish (WPI Fluoro Dish, FD35–100). The dish was centrifuged 3 min, 600 g, RT and installed in the microscope incubator at 37°C (Fig. 12A) or room temperature (Fig. 12BCD).
- 2D preformed fiber-retraction assay (Fig. 12E): Preformed fluorescent fibrin fibers were prepared in the absence of platelets (2.5 μl plasma/ml PBS; fibrinogen-Alexa 488, final concentration 12.5 μg/ml; thrombin, final concentration 2.5U/ml; incubation 15 min RT in the dark). PRP was adjusted to 2.5×10^6^ platelets/ml with PBS and 1800 μl was transferred into a petri dish (WPI Fluoro Dish, FD35–100). The dish was centrifuged 3 min, 600 g at RT to allow spreading of the platelets. After centrifugation the supernatant was replaced by the preformed fibrin fiber suspension and the dish was installed in the microscope incubator at 37°C.

For all time-lapse videos, image acquisition was started immediately after adjusting the focal position and image parameters (<1 min) using a confocal laser-scanning microscope (ConfoBright or Dynascope, for microscope details see paragraph image acquisition).

### The model of fibrin winding-up

The platelet bulb was modelled as a straight cylinder (the radius R = 400 nm, the length = 1500 nm). We assumed that the internal cytoskeleton of the bulb underwent the swirling motion with the angular velocity Rate_swirl as it was shown in^16^ and simultaneously moved down to the bottom of the bulb with the velocity Rate. Integrins αIIbβ3 on the bulb surface were associated with internal cytoskeletal proteins,^41^ thus, they underwent the swirling and downstream motion with the same rates.

The fibrin fiber was modelled as it was done previously^24,25^ with several simplifications. The fiber composed of N equally spaced nodes connected by elastic springs (supplementary Fig. 3A). The spring equilibrium length was L and it was equal to the initial distance between nodes. The fiber was located in the XY-plane and the angle α between fiber and x-axis was variable (Table 1). The motion of the node i in solution was described by the second Newton’s law of motion. The system was assumed to be in an overdamped regime (mä = 0):

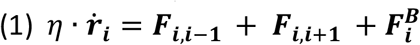

where ***r***_***i***_ was the position vector of the node i in three-dimensional coordinates (x,y,z); η was the drag coefficient for the fiber node in the blood serum; ***F***_***i***,***i***–**1**_ = *K*_*r*_ *∙* (∆***r***_*i,i*–1_ *−* ***L***) and ***F***_***i***,***i***+**1**_ = *K*_*r*_ *∙* (∆***r***_*i,i*+1_ *−* ***L***) were the vector forces which acted on the node i from the node i-1 or i+1 according to the Hook’s law due to the spring elongation or shortening (∆***r***_*i,i*–1_ and ∆***r***_*i,i*+1_ was the distance between nodes i and i-1 or the nodes i and i+1; Kr was the fiber spring constant); 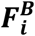 was the Brownian force which acted on the node i due to thermal fluctuations. To solve the equations for the nodes in solution, the system of equations was discretized and the Heun’s method was used. According to this method, the evolution of coordinates in time for the differential equation

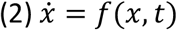

was calculated using a two-step algorithm:

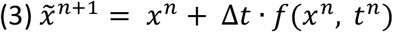

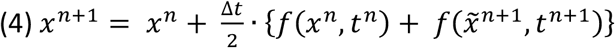

where ∆t was the time step used in calculation and discretization scheme; tn+1 = tn + ∆t; xn was coordinate at the tn moment of time.

Thus, the evolution of the node i coordinates in time was calculated according to a two-step process (in a vector form):

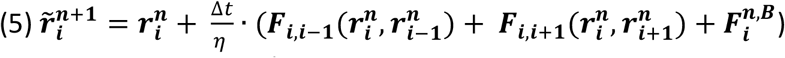

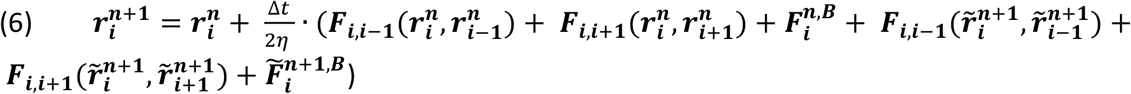

The Brownian force in three dimensions was calculated as described previously.^24^ Briefly, the mean square displacement of a particle over a single time step was 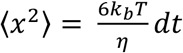, and the discretized Brownian force acting on the node i was 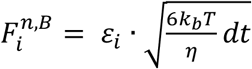, where εi was sampled from the standard normal distribution.

At the initial time point, the node 0 was bound to the integrin receptor on the bulb surface (top view, supplementary Fig. 3B). The distance of the binding point from the bulb top was D. The height of the binding point from the bulb surface was equal to the integrin αIIbβ3 molecule length (H = 20 nm) in its elongated conformation.^42^ The equations for the node 0 swirling and downstream motion were:

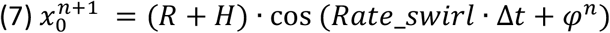

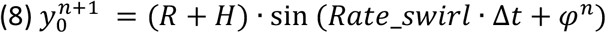

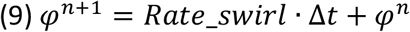

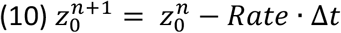

When the node i entered the light-magenta zone (supplementary Fig. 3B), and its distance from the bulb surface became smaller than H, it bound the integrin molecule and its equations of motion became similar to the node 0 equations (7) – (10).

The integrin αIIbβ3 surface density on the bulb surface was relatively high (the distance between the neighboring molecules was approximately 15 nm). Given such a high density, we assumed that when the node became close enough to the bulb, it immediately encountered the integrin molecule and bound it. Thus, when the distance between the node i and the bulb surface became smaller than H, node i bound to the integrin molecule and the node’s equations of motion became similar to the node 0 equations (7) – (10). The binding was assumed to be irreversible.

The model parameters were listed in the Table 1 below. The model equations were solved using Python 3.11.

## Acknowledgement

We are grateful to Jacques Mazzega of the microscopy platform and to Arnold Fertin for image analysis. We gratefully acknowledge the MicroCell facility (GIS IBiSA, ISdV, IAB) for its technical assistance and the microscopy system. The MicroCell facility is a member of the national infrastructure France-BioImaging (https://france-bioimaging.org/). We thank Rémy Sadoul, Sandrine Fraboulet and Alexander Bershadsky for critical reading of the manuscript. This work was supported by the Fondation Recherche Médicale (FRM, grant number DEI20151234416) and the University Grenoble Alpes (UGA, grant number AGIR-POLE FRAG15CS08).

## Declaration of Interest statement

The authors declare that they have no competing financial interests.

## Authorship contributions

A.G. developed the microscope with adaptive optics corrections and acquired images T.K. and M.P. established the model and simulations. A.-S.R. performed experiments, F.A. helped with image analyses and rendering. A.E. and J.-Y.R. acquired TEM images. L.L. participated in manuscript writing. K.S. initiated and planned the study, performed experiments, interpreted the results and wrote the manuscript. All authors read the manuscript and agreed to its content.

## Data sharing statement

All raw data are stored on the server of our microscope platform and can be accessed upon request.

## Notes

### Competing Interest Statement

The authors have declared no competing interest.

### Summary of Updates

We have revised the manuscript in accordance to the reviewer's comments. Several parts of the manuscript, have been extensively rewritten to add explanations and clarify our hypothesis and claims, this also led us to add four new references. In addition, we have added the following new figures: - new part of figure 2. Figure 2E shows a platelet in a constrained clot after 4h of retraction with the fibrin cage around the platelet center still present. The actin staining of the platelet shows that radial actin fibers are present in each bulb extending to the platelet center. This observation supports our hypothesis that in each bulb an individual cytoskeletal swirling could take place resulting in the accumulation of fibrin fibers at the base of each bulb. - modification of figure 3, to include the criteria used to define four categories of platelets and associated fibers in the 2D fiber retraction assay (new Fig. 3C). - new figure 13, illustrating the quantification of fibrin fiber compactions mediated by platelets in the 2D fiber retraction assay and the rotational movement of a fiber mass (using time-lapse sequences compiled in video 9). - new supplementary figure 1, showing the result of a new model simulation in the absence of cytoskeleton swirling. Under this condition the fibrin fiber does not loop around the platelet bulb.

## References

1. Litvinov RI, Weisel JW. Fibrin mechanical properties and their structural origins. Matrix Biol. 2017;60–61:110–123.

2. Tutwiler V, Maksudov F, Litvinov RI, Weisel JW, Barsegov V. Strength and deformability of fibrin clots: Biomechanics, thermodynamics, and mechanisms of rupture. Acta Biomater. 2021;131:355–369.

3. Adair BD, Alonso JL, van Agthoven J, et al. Structure-guided design of pure orthosteric inhibitors of alphaIIbbeta3 that prevent thrombosis but preserve hemostasis. Nat Commun. 2020;11(1):398.

4. Litvinov RI, Weisel JW. Blood clot contraction: Mechanisms, pathophysiology, and disease.Res Pract Thromb Haemost. 2023;7(1):100023.

5. Peshkova AD, Weisel JW, Litvinov RI. A novel technique to quantify the kinetics of blood clot contraction based on the expulsion of fluorescently labeled albumin into serum. J Thromb Haemost. 2024;22(6):1742–1748.

6. Tucker KL, Sage T, Gibbins JM. Clot Retraction. In: Gibbins J., Mahaut-Smith M. (eds) Platelets and Megakaryocytes. Methods in Molecular Biology (Methods and Protocols). 2012;788.

7. Sun Y, Oshinowo O, Myers DR, Lam WA, Alexeev A. Resolving the missing link between single platelet force and clot contractile force. iScience. 2022;25(1):103690.

8. Peshkova AD, Rednikova EK, Khismatullin RR, et al. Red blood cell aggregation within a blood clot causes platelet-independent clot shrinkage. Blood Adv. 2025.

9. Lam WA, Chaudhuri O, Crow A, et al. Mechanics and contraction dynamics of single platelets and implications for clot stiffening. Nat Mater. 2011;10(1):61–66.

10. Kim OV, Litvinov RI, Alber MS, Weisel JW. Quantitative structural mechanobiology of platelet-driven blood clot contraction. Nat Commun. 2017;8(1):1274.

11. Leistikow EA, Barnhart MI, Escolar G, White JG. Receptor–ligand complexes are cleared to the open canalicular system of surface-activated platelets. British Journal of Haematology. 1990;74(1):93–100.

12. Ramanujam RK, Lavi Y, Poole LG, Bassani JL, Tutwiler V. Understanding blood clot mechanical stability: the role of factor XIIIa-mediated fibrin crosslinking in rupture resistance. Res Pract Thromb Haemost. 2025;9(4):102871.

13. Spiewak R, Gosselin A, Merinov D, et al. Biomechanical origins of inherent tension in fibrin networks. J Mech Behav Biomed Mater. 2022;2022:105328.

14. Poulter NS, Pollitt AY, Davies A, et al. Platelet actin nodules are podosome-like structures dependent on Wiskott-Aldrich syndrome protein and ARP2/3 complex. Nat Commun. 2015;2015:7254.

15. Joubert C, Grichine A, Dolega M, et al. Spatial and temporal characterization of cytoskeletal reorganizations in adherent platelets. Platelets. 2024;35(1):2422437.

16. Tee YH, Shemesh T, Thiagarajan V, et al. Cellular chirality arising from the self-organization of the actin cytoskeleton. Nat Cell Biol. 2015;17(4):445–457.

17. Tamada A, Igarashi M. Revealing chiral cell motility by 3D Riesz transform-differential interference contrast microscopy and computational kinematic analysis. Nat Commun. 2017;8(1):2194.

18. Tamada A, Kawase S, Murakami F, Kamiguchi H. Autonomous right-screw rotation of growth cone filopodia drives neurite turning. J Cell Biol. 2010;188(3):429–441.

19. Gaertner F, Ahmad Z, Rosenberger G, et al. Migrating Platelets Are Mechano-scavengers that Collect and Bundle Bacteria. Cell. 2017;171(6):1368–1382 e1323.

20. Estévez-Torres A, Baigl D. DNA compaction: fundamentals and applications. Soft Matter. 2011;2011:6746.

21. Brzoska T, Suzuki Y, Mogami H, Sano H, Urano T. Binding of thrombin-activated platelets to a fibrin scaffold through α(IIb)β3 evokes phosphatidylserine exposure on their cell surface. PloS one. 2013;8(2):e55466–e55466.

22. Tutwiler V, Litvinov RI, Lozhkin AP, et al. Kinetics and mechanics of clot contraction are governed by the molecular and cellular composition of the blood. Blood. 2016;127(1):149–159.

23. Leon C, Eckly A, Hechler B, et al. Megakaryocyte-restricted MYH9 inactivation dramatically affects hemostasis while preserving platelet aggregation and secretion. Blood. 2007;110(9):3183–3191.

24. Michael C, Pancaldi F, Britton S, et al. Combined computational modeling and experimental study of the biomechanical mechanisms of platelet-driven contraction of fibrin clots. Commun Biol. 2023;6(1):869.

25. Kliuchnikov E, Peshkova AD, Vo MQ, et al. Exploring effects of platelet contractility on the kinetics, thermodynamics, and mechanisms of fibrin clot contraction. NPJ Biol Phys Mech. 2025;2(1):6.

26. Jansen KA, Zhmurov A, Vos BE, et al. Molecular packing structure of fibrin fibers resolved by X-ray scattering and molecular modeling. Soft Matter. 2020;16(35):8272–8283.

27. Hagmann J. Pattern formation and handedness in the cytoskeleton of human platelets.Proc Natl Acad Sci U S A. 1993;90(8):3280–3283.

28. Morgenstern E, Ruf A, Patscheke H. Ultrastructure of the interaction between human platelets and polymerizing fibrin within the first minutes of clot formation. Blood Coagul Fibrinolysis. 1990;1(4–5):543–546.

29. White JG, Krivit W, Vernier RL. The Platelet-Fibrin Relationship in Human Blood Clots: An Ultrastructural Study Utilizing Ferritin-Conjugated Anti-Human Fibrinogen Antibody. Blood. 1965;25:241–257.

30. Cohen I, Gerrard JM, White JG. Ultrastructure of clots during isometric contraction. J Cell Biol. 1982;93(3):775–787.

31. Morgenstern E, Daub M, Dierichs R. A new model for in vitro clot formation that considers the mode of the fibrin(ogen) contacts to platelets and the arrangement of the platelet cytoskeleton. Ann N Y Acad Sci. 2001;936:449–455.

32. Kim OV, Litvinov RI, Gagne AL, French DL, Brass LF, Weisel JW. Megakaryocyte-induced contraction of plasma clots: cellular mechanisms and structural mechanobiology. Blood. 2024;143(6):548–560.

33. Williams EK, Oshinowo O, Ravindran A, Lam WA, Myers DR. Feeling the Force: Measurements of Platelet Contraction and Their Diagnostic Implications. Semin Thromb Hemost. 2019;45(3):285–296.

34. Myers DR, Qiu Y, Fay ME, et al. Single-platelet nanomechanics measured by high-throughput cytometry. Nat Mater. 2017;16(2):230–235.

35. Mercey O, Reymond L, Lemaître F, et al. HAK-actin, U-ExM-compatible probe to image the actin cytoskeleton. bioRxiv. 2025:2025.2008.2026.672318.

36. Kovalenko TA, Giraud MN, Eckly A, et al. Asymmetrical Forces Dictate the Distribution and Morphology of Platelets in Blood Clots. Cells. 2021;10(3).

37. Grichine A, Jacob S, Eckly A, et al. The fate of mitochondria during platelet activation. Blood Adv. 2023;7(20):6290–6302.

38. Gambarotto D, Hamel V, Guichard P. Ultrastructure expansion microscopy (U-ExM). Methods Cell Biol. 2021;161:57–81.

39. Schindelin J, Arganda-Carreras I, Frise E, et al. Fiji: an open-source platform for biological-image analysis. Nat Methods. 2012;9(7):676–682.

40. Eckly A, Strassel C, Cazenave JP, Lanza F, Leon C, Gachet C. Characterization of megakaryocyte development in the native bone marrow environment. Methods Mol Biol. 2012;788:175–192.

41. Bidone TC, Skeeters AV, Oakes PW, Voth GA. Multiscale model of integrin adhesion assembly. PLoS Comput Biol. 2019;15(6):e1007077.

42. Xu XP, Kim E, Swift M, Smith JW, Volkmann N, Hanein D. Three-Dimensional Structures of Full-Length, Membrane-Embedded Human alpha(IIb)beta(3) Integrin Complexes. Biophys J. 2016;110(4):798–809.

43. Weisel JW, Litvinov RI. Fibrin Formation, Structure and Properties. Sub-cellular biochemistry. 2017;82:405–456.

