## Supplementary material for "Winding-Up of Fibrin Fibers as a Novel Mechanism of Platelet-Mediated Fiber Compaction": suplementary figures and video legends

### Supplementary figures and legends

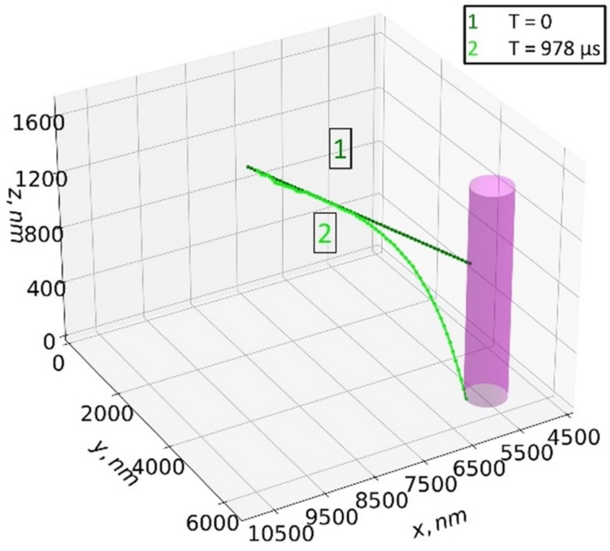


supplementary Figure 1

***Supplementary Figure 1: The model of fibrin accumulation without cytoskeleton swirling.*** *Position of the fibrin fiber at the initial (1, green) and final (2, lime) time points. After being pulled to the base of the bulb, no fibrin loop was observed. The platelet bulb is depicted as a magenta cylinder.*

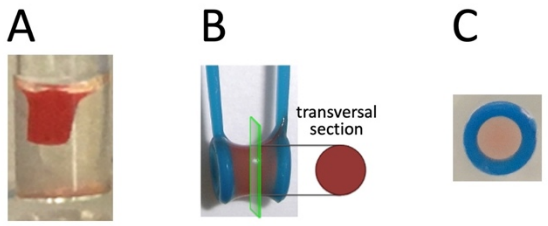


supplementary Figure 2

***Supplementary Figure 2: Illustration of clot retraction assays used in this study***
*A) Unconstrained clot retraction (for figure 1AB)*
*B) Constrained clot retraction between two holders (for figure 1C)*
*C) Constrained clot retraction within an inoculation loop (for figure 2)*

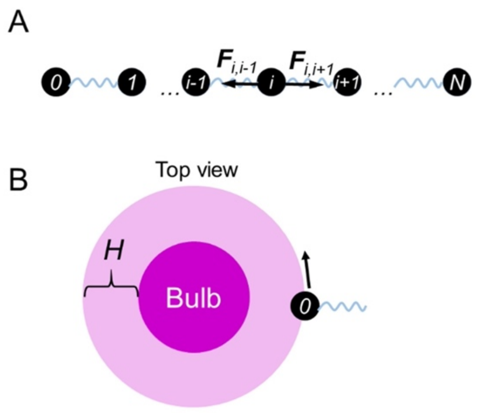


supplementary Figure 3

***Supplementary Figure 3: Model of fibrin fiber binding to the platelet bulb and fibrin winding-up.***

*A) The model of a fibrin fiber in solution. The fiber was modeled as a sequence of N nodes (black dots) which were connected by elastic springs (blue).* $F_{i,i-1}$*/* $F_{i,i+1}$ *were vector forces which acted on the node i according to the Hook's law, the black vectors show the force directions.*

*B) The system configuration at time point T=0. Magenta circle: internal volume of a bulb. Light-magenta: the binding region. If the node i entered the binding region, it bound to the integrin molecule. The black arrow shows the rotation direction of the cytoskeleton and integrin molecules.*

### Supplemental Videos (Time-lapse and Animations)

**Video 1 (associated to figure 1B)**

*Depth color-coded time-lapse of an unconstrained clot retraction (1x$10^8$ platelets per ml in 50% plasma/50% PBS) in presence of fibrinogen-Alexa 488. Fibrin fibers (green) present in z-projections of each time point are also shown upper right corner. Acquisition was started immediately after thrombin addition at 100 $\mu$m above the bottom of the well and image stacks were acquired (61 focal planes, step size 0.5 $\mu$m) for a time period of 20:27 min (100 frames with a time interval of 12.5 seconds, scale bar 10 $\mu$m).*

**Video 2 (associated to figure 2)**

*Two typical examples of platelets in a constrained clot with attached fibrin fibers (green) are shown (1x$10^7$ platelets per ml PBS/50% plasma and fibrinogen-Alexa 488). Clot retraction was allowed to take place for 15 min before fixation and immunofluorescence staining of the $\alpha$IIb integrin subunit (magenta). Samples were then processed for expansion (scale bars 10 $\mu$m = 2.5 $\mu$m after correction for expansion). Left panels, scrolling through the image stack; right panels, 3D reconstruction.*

*The lower panel shows a 3D reconstruction of a typical example of a platelet in a constrained clot (1x$10^7$ platelets per ml PBS/50% plasma, retraction 4h) stained for the actin cytoskeleton (magenta) and then processed for expansion. The sample was then stained for total proteins (platelet proteins as well as fibrin fibers) using the* NHS-Ester DyLight 650 (cyan; *scale bar 10 $\mu$m = 2.5 $\mu$m after correction for expansion)*

**Video 3 (associated to figure 4)**

*Two typical examples of fiber accumulations (green) above spread platelets in the 2D fiber-retraction assay (4 $\mu$l plasma per ml PBS, see methods section) are shown. Samples were stained for the $\alpha$IIb integrin subunit (magenta) and expanded (scale bars 10 $\mu$m = 2.5 $\mu$m after correction for expansion). Upper panels, scrolling through the image stack; lower panels, 3D reconstruction.*

**Video 4 (associated to figure 5)**

*Two examples of platelets with bulbs encircled by fibrin fibers (green) in the 2D fiber-retraction assay (4 $\mu$l plasma/ml, see methods section) are shown. Samples were stained for the $\alpha$IIb integrin subunit (A, B; magenta) or the myosin light chain (MLC; C,D; magenta) and processed for expansion (scale bars 10 $\mu$m = 2.5 $\mu$m after correction for expansion). Upper panels, scrolling through the image stack; lower panels, 3D reconstruction.*

**Video 5 (associated to figure 6)**

*Two examples of platelets with twisted fibrin fibers (green) are shown in the 2D fiber-retraction assay (4 $\mu$l plasma/ml, see methods section). Samples were stained for myosin II (A; magenta) or phosphorylated myosin light chain (p-MLC; C,D; magenta) and processed for expansion (scale bars 10 $\mu$m = 2.5 $\mu$m after correction for expansion). Upper panel 3D reconstruction (Fig. 6A). Middle panel, scrolling through the image stack (Fig. 6C); lower panel, 3D reconstruction (Fig. 6D).*

*Please note also the strong fiber accumulation around the platelets similar to platelets shown in video 4/figure 5.*

**Video 6 (associated to figure 7)**

*Four examples of a fibrin rosette associated with spread platelets in the 2D fiber-retraction assay (7 $\mu$l plasma per ml PBS, see methods section) are shown. Samples were stained for the $\alpha$IIb integrin subunit as well as for actin and processed for expansion (scale bars 10 $\mu$m = 2.5 $\mu$m after correction for expansion). 3D image reconstructions of image stacks. Fibrin fibers (green) and integrin or actin staining (upper and lower panels respectively, magenta).*

**Video 7 (associated to figure 8)**

*Three examples of fibrin (green) and myosin (magenta) localization for spread platelets in the 2D fiber-retraction assay (4 $\mu$l plasma/ml for A, B, E, F or 7 $\mu$l plasma/ml C, D; see methods section). Samples were stained for myosin (A, B, C, D) or phosphorylated myosin light chain (p-MLC; E, F) and expanded (scale bars 10 $\mu$m = 2.5 $\mu$m after correction for expansion). Upper panels, scrolling through the image stack; lower panels, 3D reconstruction.*

**Video 8 (associated to figure 10)**

*Simulation of fiber winding-up around a platelet bulb. Angle between the fiber and the x-axis $\alpha = \pi/2.5$*

**Video 9 (associated to figure 12)**

*Live imaging of spread platelets (transmission gray) and fluorescent fibrin fibers (green), scale bars 10 $\mu$m.*

*Upper three panels : Spread platelets from three different donors are shown, which reorganize and compact fluorescent fibrin fibers.*

*Lower panel : Preformed fluorescent fibrin fibers were added to spread platelets. Culture surface coating is observed and a dense fibrin mass in the platelet center is rotating in several platelets.*
